# Redox-modulated bacterial deubiquitinase ElaD: Target recognition and suppression of K63-linked polyubiquitin accumulation in yeast

**DOI:** 10.64898/2026.06.26.730077

**Authors:** Lubhanshi Garg, Aditya Shrivastava, Gessica C. Barros, Gustavo M. Silva, Sri Rama Koti Ainavarapu

## Abstract

Bacterial deubiquitinases (DUBs) are important virulence effectors that manipulate host ubiquitin signaling during infection. ElaD, a CE-clan DUB expressed by enterohemorrhagic *Escherichia coli*, preferentially cleaves K63-linked ubiquitin chains, yet its effects on conserved cellular stress responses remain poorly understood. We demonstrate that ElaD exhibits redox-dependent DUB activity *in vitro*. In addition, we identified the molecular basis underlying the selective recognition of substrate proteins, ubiquitin and NEDD8 by ElaD. Structural and mutational analyses reveal that, beyond the conserved catalytic site, ElaD engages ubiquitin through a combination of electrostatic and hydrophobic interactions. Using Saccharomyces cerevisiae as a heterologous model system, we show that wild-type ElaD rescues the proteotoxic stress phenotype of ubp2Δ yeast cells, whereas specific ElaD mutants fail to confer a similar response. Furthermore, expression of ElaD suppresses oxidative stress–induced accumulation of K63-linked polyubiquitin and may perturb stress-associated translational regulation linked to K63 ubiquitin signaling. Consequently, cells expressing ElaD exhibit altered stress adaptation and diminished fitness during prolonged oxidative stress. Collectively, these findings indicate that ElaD perturbs ubiquitin-mediated stress signaling by counteracting K63-linked ubiquitination events that support adaptive cellular responses. Our study highlights how a bacterial DUB can reprogram conserved ubiquitin-dependent pathways and exploit host ubiquitin signaling networks to modulate cellular stress responses and protein homeostasis. These findings further suggest potential host targets of bacterial DUBs during infection.

## INTRODUCTION

Maintenance of cellular proteostasis is essential for eukaryotic viability and is tightly regulated by the ubiquitin–proteasome system (UPS), which controls protein turnover, localization, and activity [1–5]. Ubiquitination is a reversible post-translational modification in which the 76–amino-acid protein, ubiquitin, is most commonly conjugated to lysine residues on substrate proteins [6], through an enzymatic cascade involving E1 activating enzymes, E2 conjugating enzymes, and E3 ligases [2,7,8]. Ubiquitin can also be attached, albeit less frequently, to non-lysine residues such as cysteine, serine, threonine, and the substrate N-terminus [9–11]. The assembly of polyubiquitin chains broadens the signaling capacity of the UPS, as distinct linkage types encode specific cellular outcomes [12,13]. Lysine-48-linked chains primarily target substrates for proteasomal degradation [14], whereas lysine-63-linked chains function largely in non-proteolytic processes, acting as scaffolds for signaling complexes involved in DNA repair [15], kinase activation [16,17], vesicular trafficking [18,19], and cellular stress responses [20,21]. The ubiquitin signaling pathway preserves protein homeostasis by employing deubiquitylating enzymes (DUBs) to remove ubiquitin from substrates, which also maintains the intracellular free ubiquitin pool [6,22]. Although ubiquitin signaling is a defining feature of eukaryotic cells, DUBs have also been identified across diverse domains of life [23,24].

Humans encode approximately 100 DUBs [25,26] that can cleave either mono– or polyubiquitin-linked chains from the substrate. These are catalytically divided into two broad categories: cysteine proteases and metalloproteases. Majority of the DUBs are cysteine proteases and are predominantly found in eukaryotes [27]. DUBs are regulated by different post-translational modifications [28,29], substrate binding [30] and redox regulation [26,31]. Redox modulation of catalytic activity has been previously reported for several eukaryotic cysteine-based DUBs and is known to play an important role in regulating cellular processes [32,33]. Cysteine oxidation is demonstrated for various eukaryotic DUBs, which leads to reversible loss of activity. Some examples include human USP1, which is responsible for DNA damage repair, and its oxidation leads to high levels of PCNA ubiquitination [34], along with USP19 [35], USP28 [36], and CYLD, which inhibits NF-kB activation by reversing the ubiquitination of TRAF2 and TRAF6 [37–40]. Despite lacking endogenous ubiquitin pathways, several pathogens, including bacteria [41–44] and viruses [45], also encode DUBs that can act on host ubiquitin pathways. In particular, bacterial pathogens secrete effector proteins [44,46] that mimic eukaryotic ubiquitin-modifying enzymes, particularly DUBs [43] into host cells. Elastase D (ElaD), a DUB from *E. coli*, belongs to the CE-clan of cysteine proteases (MEROPS database), having a papain-fold and a conserved catalytic core with Cys at the active site [47]. During infection, host immune system responses generate oxidizing environments [48,49] for pathogen effector proteins. Despite this, redox regulation of bacterial DUBs has not yet been investigated.

Previous studies have shown that DUBs selectively recognize distinct ubiquitin linkages to regulate downstream signaling pathways [29]. The specific ubiquitin linkage type is also important in the context of bacterial DUBs that include ElaD homologs such as SseL, and XopD (Figure S1 shows structural overlap with conserved active site). SseL preferentially recognizes K63-linked polyubiquitin chains in vitro. Cellular studies involving SseL show that DUB activity helps bacterial survival inside the host [50]. ChlaDUB1, another ElaD homolog, specific to K63-polyUb, is also shown to modulate the immune response pathways, such as the NF-kB pathway for immune activation [51]. ElaD has been studied to selectively cleave K63-linked polyubiquitin chains in vitro [52]. ElaD is also expressed by enterohemorrhagic *E. coli* (EHEC), a clinically important pathogen associated with severe disease outcomes such as hemolytic uremic syndrome [42]. Together, these observations raise the possibility that ElaD could be modulating host cellular pathways. Despite increasing interest in host–pathogen ubiquitin signaling, limited studies have explored the repertoire and physiological roles of ubiquitin-modifying enzymes in *E. coli* compared to widely studied pathogens such as Salmonella [41]. Although ElaD is broadly conserved [53] little is known about the role of specific K63-linked polyubiquitin recognition by ElaD in the context of its functional impact on host ubiquitin-dependent pathways. Another aspect of substrate recognition involves ubiquitin-like proteins (Ubls) that have been identified to share the core β-grasp fold with Ub, which include SUMO, NEDD8, and ISG15 proteins [54]. Selective recognition of substrate proteins by DUBs can regulate the cascading pathways involved [55]. ElaD has been shown to selectively recognize Ub over Ubls, similar to human DUB, USP2 (selective to Ub), and SENP8 (selective to NEDD8). SENP8, a structural homolog of ElaD specific to NEDD8, discriminates substrates through interactions with specific residues [56,57]. The molecular basis of the selective recognition of Ub over Ubls by ElaD is unknown. The structural features responsible for the specific cleavage of K63-linked polyubiquitin are also yet to be investigated.

Given the prominent role of K63-linked ubiquitin signaling in stress-responsive pathways, model systems that enable studies on ubiquitin dynamics can provide valuable insight into bacterial DUB function. The budding yeast *Saccharomyces cerevisiae* has long served as a powerful model for dissecting ubiquitin biology, as the UPS is highly conserved across eukaryotes [58–60]. Importantly, yeast relies extensively on K63-linked polyubiquitination as a regulatory signal during oxidative [20] and proteotoxic stress, where these chains control translation [61], autophagy [62], and protein quality surveillance [63]. More broadly, K63-linked ubiquitin signaling in yeast under oxidative stress functions as a regulatory signal, conceptually analogous to the roles of K63-linked ubiquitin chains in mammalian stress and immune signaling pathways, including NF-κB activation [16,59,64]. Previous studies have demonstrated that expression of bacterial effectors in yeast can faithfully recapitulate key aspects of host-pathogen interactions, including modulation of translation and cell growth [58,60,65–67]. Whether bacterial DUBs alter ubiquitin accumulation during oxidative stress conditions has not been explored.

In this study, we exploit this conserved cellular framework to establish a yeast-based system for characterizing the activity, specificity, and physiological impact of the bacterial deubiquitinase ElaD. Here, we have characterized ElaD as a redox-modulated bacterial DUB and elucidated the molecular basis of substrate recognition. We also demonstrate that selectivity can be reprogrammed through targeted mutation in substrate proteins. Moreover, ElaD suppresses the accumulation of K63-linked polyubiquitin chains under oxidative stress in yeast, and its DUB activity rescues yeast from proteotoxic stress.

## RESULTS

### DUB activity of ElaD is redox modulated

Eukaryotic cysteine-based DUBs have been reported to be redox modulated and also regulate various cellular pathways [32]. However, the redox modulation of cysteine proteases in bacteria remains to be studied. ElaD is one of the CE-clan DUBs that specifically cleaves K63-linked polyubiquitin chains [52]. We examined whether ElaD activity is regulated by redox conditions. His-tagged ElaD was expressed in *E. coli* and purified using immobilized metal affinity chromatography as described in the Methods section. Figure 1A shows the scheme for monitoring the DUB activity of ElaD using a fluorescently labelled protein, Ub-Rhodamine, as a substrate [68]. An increase in fluorescence is found to be ElaD concentration dependent, demonstrating that ElaD cleaves ubiquitin from the Ub-Rho conjugate. On mutating the catalytic Cys to Ala (ElaD C317A), the DUB activity was abolished, as shown in Figure 1B. Further, ElaD exhibits enhanced DUB activity under reducing conditions, while decreased activity under oxidizing conditions. To elucidate the redox dependence of ElaD activity, the enzyme was pretreated with either hydrogen peroxide (H_2_O_2_) or dithiothreitol (DTT) prior to substrate addition. ElaD activity decreased progressively with increasing concentrations of H_2_O_2_, whereas inhibition was reversed upon treatment with DTT (Figure 1C), indicating reversible redox-regulation. Notably, ElaD displays redox-sensitive behavior similar to eukaryotic DUBs Ubp2 [63] and USP1 [33], both of which are regulated through reversible oxidation. Despite the limited structural similarity of ElaD with the above-mentioned DUBs, the functional convergence suggests that redox sensitivity may represent a conserved regulatory feature of cysteine DUBs. Together, these findings raise the possibility that redox control of DUB activity enables dynamic control of cellular processes in response to changes in redox environment.

**Figure 1.**
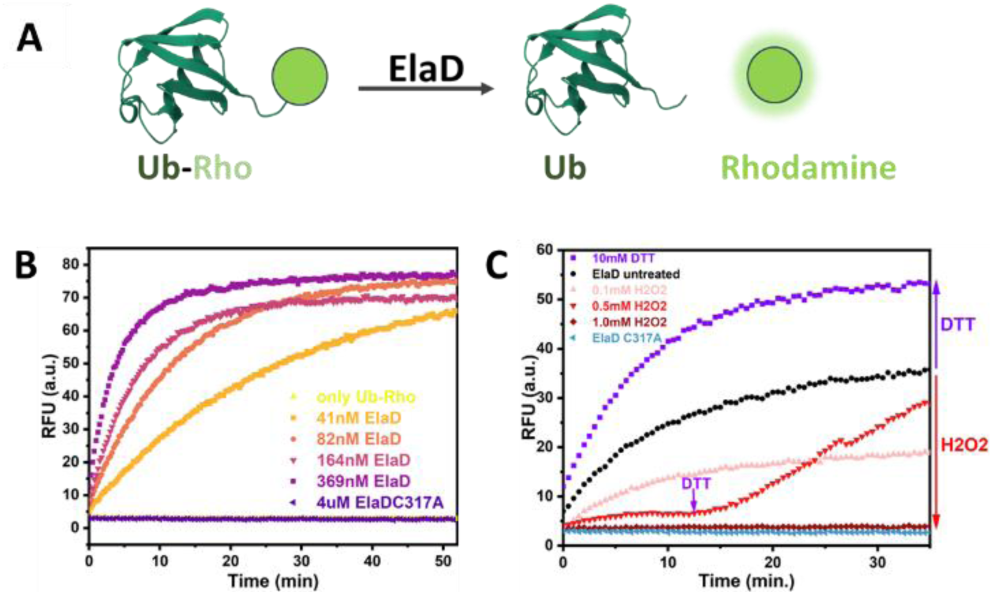
Redox-sensitive DUB activity of ElaD. A, Schematic representation of the ubiquitin cleavage assay. Ubiquitin conjugated to rhodamine 110 (Ub-Rho) was used as the substrate. Ub-Rho cleavage was monitored by recording fluorescence at 535 nm (excitation at 485 nm). B, ElaD, and its mutant were incubated with 0.75 µM Ub-Rho, and enzyme kinetics were monitored by measuring fluorescence. Substrate Ub-Rho shows no change in fluorescence over time (overlaps with ElaD C317A data). C, Enzymatic activity of ElaD (82 nM) was assessed *in vitro* using 0.75 µM Ub-Rho. Prior to substrate addition, ElaD was pre-incubated for 5 min with H_2_O_2_ (0.1 mM, 0.5 mM, or 1 mM) or with 10 mM DTT. In one condition, 10 mM DTT was added after 12 min to reduce ElaD that had been oxidized with 0.5 mM H_2_O_2_.

### Modeling the ElaD–ubiquitin complex highlights the binding site

To investigate potential interactions between ElaD and ubiquitin, we generated a structural model of ElaD-Ub complex using ColabFold (AlphaFold2-multimer v3) [69,70]. Figure 2A shows a representative complex of one of the four independently seeded models. All models converged on an identical binding mode, with maximum Cα RMSD of 0.6 Å over the interface core (pruned atoms) and 1.3–2.7 Å over all aligned atoms, indicating minimal conformational heterogeneity (Table S1). The top-ranked unrelaxed model displayed a predicted TM-score (pTM) of 0.83 and an interface TM-score (ipTM) of 0.93, consistent with a highly confident inter-chain orientation. In addition, inter-chain regions of the predicted aligned error (PAE) matrix were uniformly low (<5–10 Å), further supporting a well-defined relative positioning of the two proteins (Figure S2). The binding site of Ubq C-terminal chain with key residues labeled is shown in Figure 2B.

**Figure 2.**
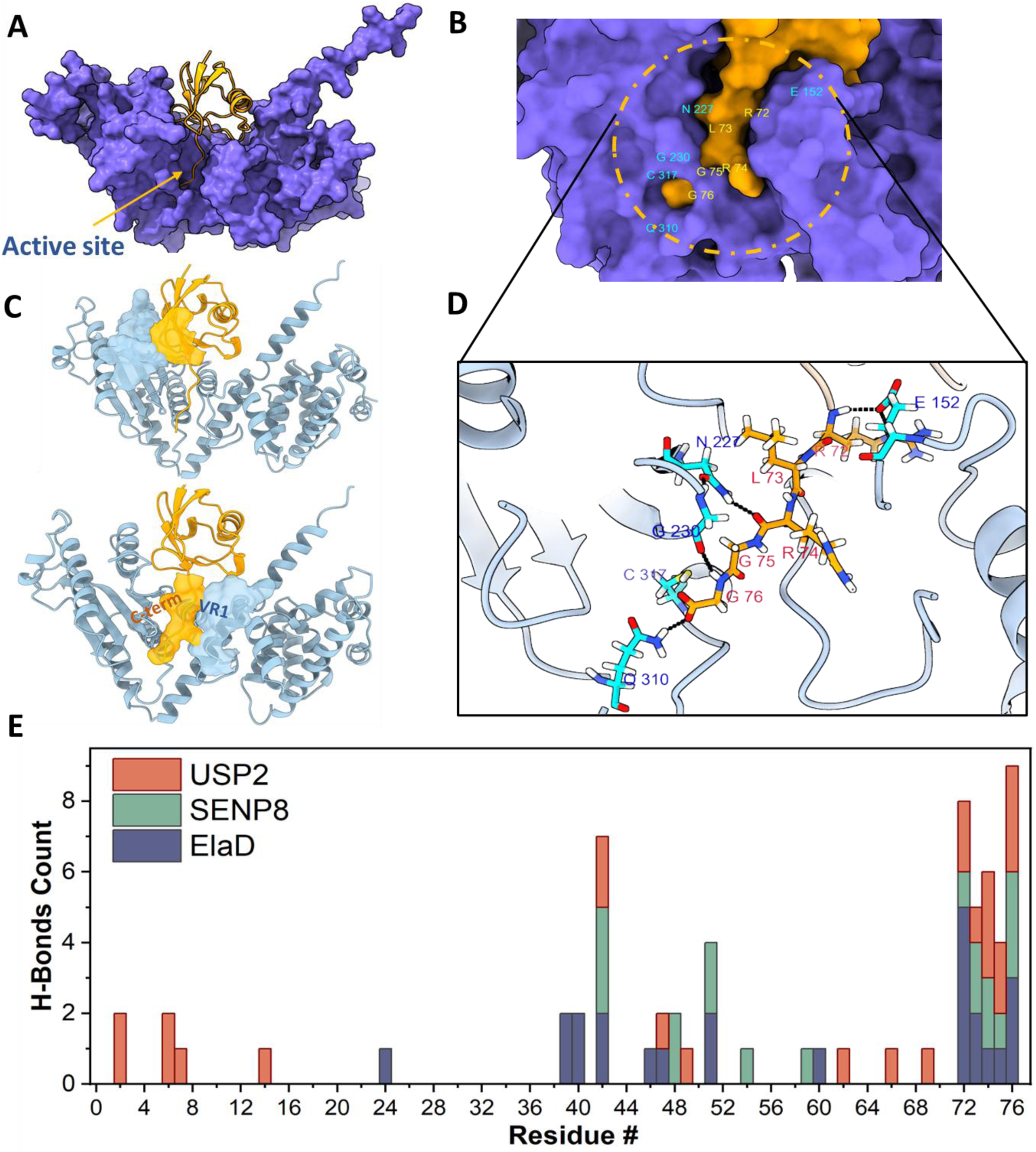
Structural analysis to identify key residues involved in substrate recognition. A, ElaD (UniprotID: Q8XCY9) in complex with Ub (PDB: 1UBQ) complex as predicted by ColabFold using AlphaFold multimer (v3). B, Surface representation showing the ubiquitin C-terminal tail interacting with the ElaD active site cleft. C, Cartoon representation showing the interaction of the ubiquitin hydrophobic patch (top), the ElaD variable region (VR1) with ubiquitin surface residues. D, Cartoon representation showing hydrogen-bond interactions among residues, visualized using RCSB ChimeraX. Hydrogen bonds are indicated as black dashed lines. E, Residue-wise distribution of H-bond interactions across the substrate sequence for three DUBs: USP2 (salmon), SENP8 (green) and ElaD (blue) The residue # corresponds to Ub for USP2/ElaD, and NEDD8 for SENP8.

Analysis of the ElaD(A)-Ub(B) interface in ChimeraX [71] revealed a buried surface area of 1608 Å^2^, comparable to that observed in characterized DUB–ubiquitin complexes [56]. The interface contains numerous polar and hydrogen-bonding contacts, including those between Glu152(A)-Arg72(B), and Lys176(A)–Glu51(B), suggesting that electrostatic interactions contribute to ElaD-ubiquitin recognition (Figure S3). In addition, canonical hydrophobic interactions involving the Ub Ile44 patch also contribute to the binding (top panel in Figure 2C). Furthermore, the bottom panel in Figure 2C highlights surface contacts between ElaD variable-region1 (aa 147-157) and the ubiquitin C-terminal tail, positioning ubiquitin for catalysis. Figure 2D depicts hydrogen bonds between interacting residues as dashed lines. Contact-based analysis (distance cutoff < 5 Å) identified 61 interface residues distributed across the catalytic cleft of ElaD and the C-terminal β-sheet and *α*-helix of ubiquitin. A list of interacting residues with corresponding hydrogen-bond distances is provided in Supplementary Table S2a. A complete list of all the contacts is provided in Table S2b.

To narrow down specific residues that form the structural basis of substrate recognition, we compared their predicted hydrogen-bond contact profiles with the experimental interfaces of two well-characterized DUBs: human USP2 bound to ubiquitin (PDB ID: 2HD5) and SENP8 bound to NEDD8 (PDB ID: 1XT9). As shown in Figure 2E, the distribution of hydrogen bonds along the substrate sequence reveals distinct binding strategies. The USP2-Ub interface is expansive, utilizing contacts spread across the entire ubiquitin sequence. In contrast, the interactions for both SENP8 and ElaD are concentrated at the C-terminal tail of their respective substrates. Previous studies have demonstrated that substrate selectivity can be reversed for both USP2 and SENP8. For USP2, this discrimination is governed by the substrate N-terminal residues [Ub residues 4, 12, 14, and 72] [72], whereas SENP8 relies on a distinct set of determinants [NEDD8 N51/A72] (49). The list of H-bonds at the USP2-ubiquitin and SENP8-NEDD8 interfaces is provided in Supplementary Tables S3 and S4, respectively. Structural and mutational studies of the SENP8–NEDD8 complex have established that substrate recognition is mediated by interactions involving the C-terminal tail of NEDD8, particularly residue 72, which differs between NEDD8 (Ala72) and Ub (Arg72). Consistent with this, SENP8 has been reported to recognize a ubiquitin double mutant (Ub E51N/R72A), underscoring the importance of these positions in substrate discrimination. Because our interaction analysis indicates that ElaD lacks significant contact with the N-terminal region of ubiquitin, we hypothesized that its selectivity mechanism more closely resembles that of SENP8. Consequently, we targeted substrate residues Glu51 and Arg72 for mutational analysis to determine whether the substrate preference of ElaD could be fundamentally reprogrammed.

### ElaD substrate selectivity is reprogrammed through targeted mutation of Glu 51 and Arg72 in ubiquitin

ColabFold modeling of the ElaD-Ub complex predicts a well-defined interface containing multiple polar/electrostatic contacts, highlighting ubiquitin residues Glu51 and Arg72 as candidate determinants of productive recognition. ElaD preferentially recognizes ubiquitin over NEDD8, but the molecular basis of selective recognition remains unsolved. We therefore asked whether modifying these residues (Ub E51 and R72) could reprogram ElaD to recognize and cleave NEDD8. NEDD8 is a ubiquitin-like modifier that shares ∼60% sequence identity [73] and a highly conserved β-grasp fold with ubiquitin. To assess substrate specificity, a fusion substrate, Snooptag-(I27)_2_-Ub-NEDD8-Spytag-His_6_ (Figure 3A) was constructed. The construct was designed so that cleavage products could be directly resolved by SDS-PAGE based on differences in the molecular mass of cleaved protein fragments. For simplicity, we refer to this substrate protein as Ub-NEDD8. Figure 3B shows sequence alignment and structural overlap of Ub and NEDD8. The identities of the mutant substrate variants were confirmed by ESI-MS analysis of the purified proteins (Figure S4). Figure 3C highlights the predicted interacting residues in ElaD (Glu152 and Lys176), consistent with electrostatic contacts playing a role in substrate recognition. DUB activity of ElaD was tested by incubating Ub-NEDD8 wild-type (WT) vs mutant substrates with Silk–ElaD fusion protein. SDS–PAGE analysis revealed efficient cleavage of the Ub-NEDD8 WT substrate at the ubiquitin C-terminus within 30 minutes (Figure 3D). Reciprocal substitution at position 51(Ub E51N) had a limited effect on this cleavage pattern at the 30 min endpoint. In contrast, substitution at position 72 (Ub R72A) strongly reduced cleavage at ubiquitin C-terminal, without detectable cleavage at NEDD8 C-terminus. Interestingly, when both positions were exchanged in Ub-NEDD8 51/72, ElaD cleaved at the NEDD8 C-terminus. Figure 3D shows a new band consistent with the release of the C-terminal Spytag-His_6_ fragment. The band corresponds to a mass of ∼9 kDa on the gel, due to its highly basic residues (pI∼10). Similar enzymatic assays were performed with His_6_-ElaD for 30 min endpoint and 4 hr endpoint with similar results as in the case of Silk-ElaD (Figure S5). Together, these data indicate that residues E51 and R72 act as key determinants that bias ElaD toward ubiquitin, with residue 72 exerting a stronger effect, while the combined substitutions reprogram substrate selectivity toward NEDD8.

**Figure 3.**
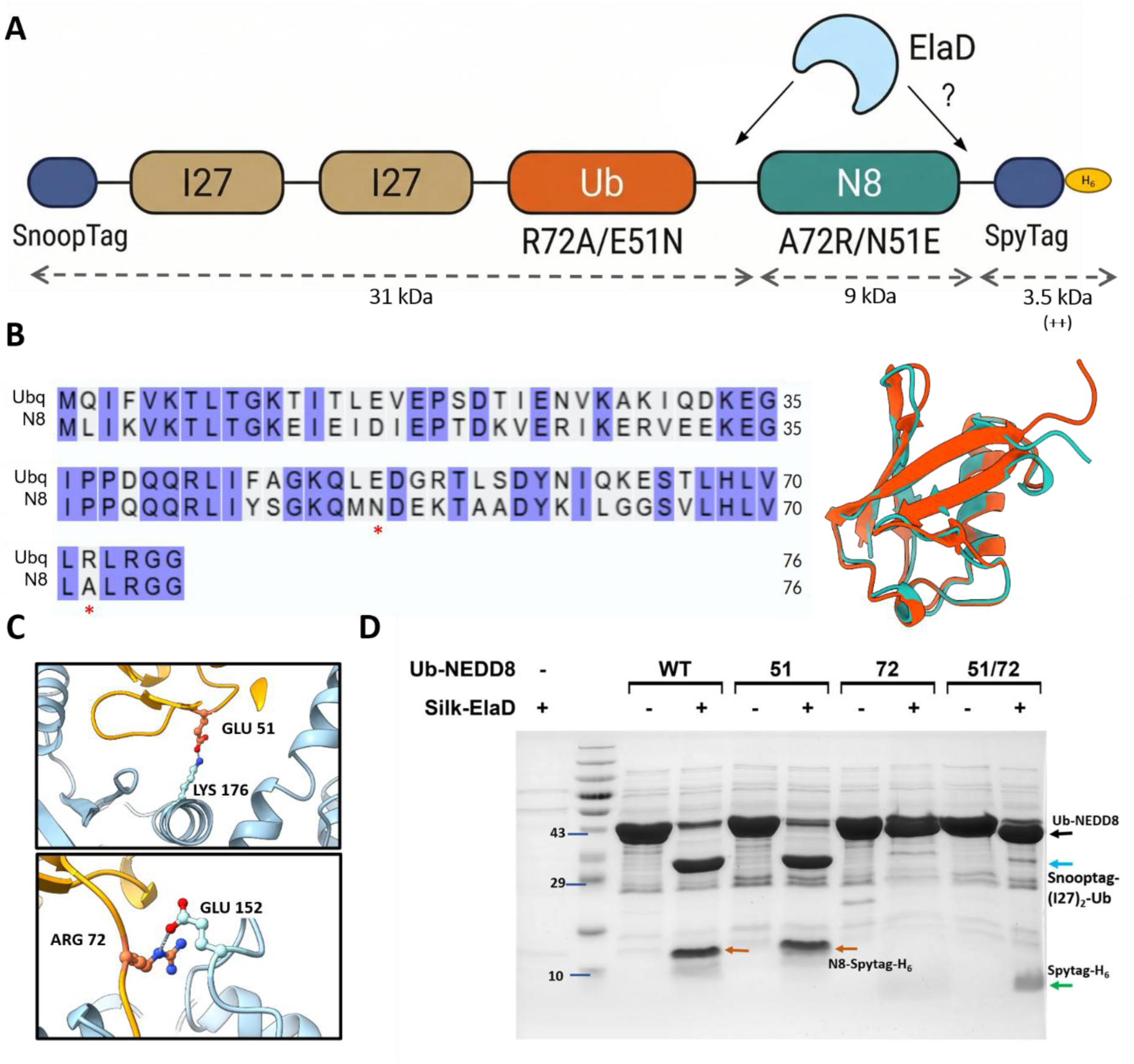
Mutational analysis confirms molecular determinants of specificity. A, Schematic for the substrate construct Snooptag-(I27)_2_-Ub-N8-Spytag. B, Structural overlap in ChimeraX of Ub (PDB:1UBQ) with NEDD8 (PDB:1NDD). Sequence alignment highlights conserved residues and key residues for interaction. C, Cartoon representation of predicted interaction (ColabFold) between specific residues in ElaD and Ub. D, Glu51 and Arg72 in Ub are critical for recognition; enzymatic cleavage assays followed by gel electrophoresis show that mutant proteins are not efficiently recognized by ElaD.

### The kinetics of substrate cleavage is altered by specific mutations

Time-course analysis corroborated the endpoint cleavage patterns, showing rapid processing of the Ub-NEDD8 WT and Ub-NEDD8 51 mutant substrate, while the Ub-NEDD8 72 mutant remained cleavable but with significantly reduced kinetics, indicating that substitution at position 72 (Ub R72A) slows ElaD-mediated cleavage. In contrast, Ub-NEDD8 51/72 double mutant displayed diminished accumulation of the major ubiquitin cleavage product and instead produced a prominent low-molecular-weight fragment consistent with preferential cleavage at the NEDD8 C-terminus (Figure 4A). Uncropped SDS-PAGE gel data is provided in Figure S6. Another 4-hour endpoint assay confirmed that both Ub-NEDD8 72 and Ub-NEDD8 51/72 remain cleavable, with increased ubiquitin-junction processing at later time points (Figure S7), indicating that these substitutions alter and redirect cleavage rather than abolish it, consistent with a shift in substrate selectivity rather than absolute specificity.

**Figure 4.**
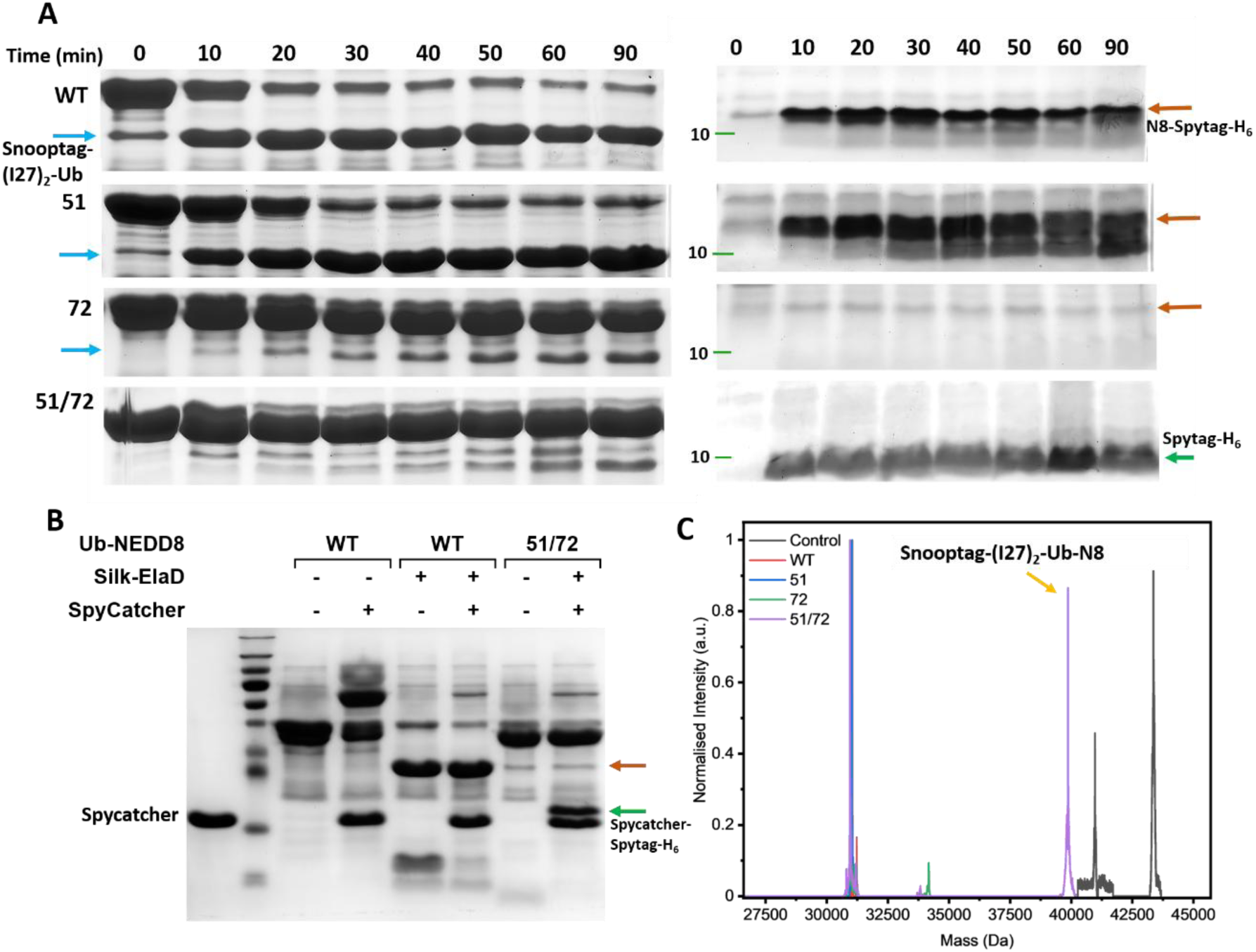
Kinetics of cleavage in different mutants and validation of cleavage at NEDD8 C-terminus in the double mutant. A, Time-course SDS–PAGE analysis of cleavage reactions for Ub-NEDD8 WT, 51, 72, and 51/72 substrates. The left panel shows the higher-molecular-weight region (∼50–30 kDa), highlighting conversion of the full-length substrate to the major cleavage product. The right panel shows the low-molecular-weight region (<15 kDa). Ub-NEDD8 WT and 51 display similar cleavage kinetics, with prominent product formation within 10 min. The Ub-NEDD8 72 mutant is cleaved slowly compared to WT, whereas the Ub-NEDD8 51/72 double mutant shows accumulation of a prominent low-molecular-mass product consistent with cleavage at NEDD8 C-terminus and release of the C-terminal Spytag-containing fragment. B, Spycatcher-Spytag-based validation of the low-molecular-mass product. Reaction products were incubated in the absence (−) or presence (+) of Spycatcher and analyzed by SDS–PAGE. In Ub-NEDD8 51/72 sample, the low molecular-weight band shows a small upward mobility shift (green arrow) upon Spycatcher addition, consistent with covalent capture of the released Spytag-containing fragment. Although the expected mass addition is small (the released Spytag-containing fragment is approximately 3.5 kDa), the shift is detectable by SDS–PAGE. In contrast, the higher-molecular-mass product (brown arrow) does not shift, indicating that it no longer retains the Spytag. C, ESI-MS analysis of different substrate proteins after incubation with ElaD. Ub-NEDD8 51/72 reaction mixture shows the peak for Snooptag-(I27)_2_-Ub-NEDD8 (yellow arrow), confirming cleavage at NEDD8 C-terminal.

The substrate protein construct included Spytag to facilitate additional validation experiments [74]. To confirm the identity of the lower mass product obtained on incubation of Ub-NEDD8 51/72 with the enzyme, reaction mixtures were incubated with Spycatcher. In the Ub-NEDD8 51/72 sample, the lower band showed an upward shift consistent with covalent capture of the released Spytag fragment, while the higher mass product remained unchanged, indicating loss of Spytag (Figure 4B). ESI-MS analysis also supported this assignment, revealing a species corresponding to the Snooptag-(I27)_2_–Ub–N8 fragment at 39864.3 Da (calculated 39864.8 Da) in Ub-NEDD8 51/72 sample (Figure 4C). Together, these results indicate that residue R72 is a key determinant for ElaD recognition of ubiquitin, and that combined substitution at positions 51 and 72 redirects cleavage toward the NEDD8 junction, highlighting their role in substrate discrimination.

### ElaD confers resistance to proteotoxic stress in yeast via specific ubiquitin recognition interfaces

To elucidate the role of ElaD in modulating host-pathogen interactions, first, we need to confirm protein activity in a biological context. To this end, we choose yeast as the model system because: 1) it has a conserved ubiquitin system similar to that of humans. 2) *Saccharomyces cerevisiae* shows increased K63-linked polyubiquitination in response to oxidative stress. Figure 5A shows the schematic for gene complementation followed by readout in yeast.

**Figure 5.**
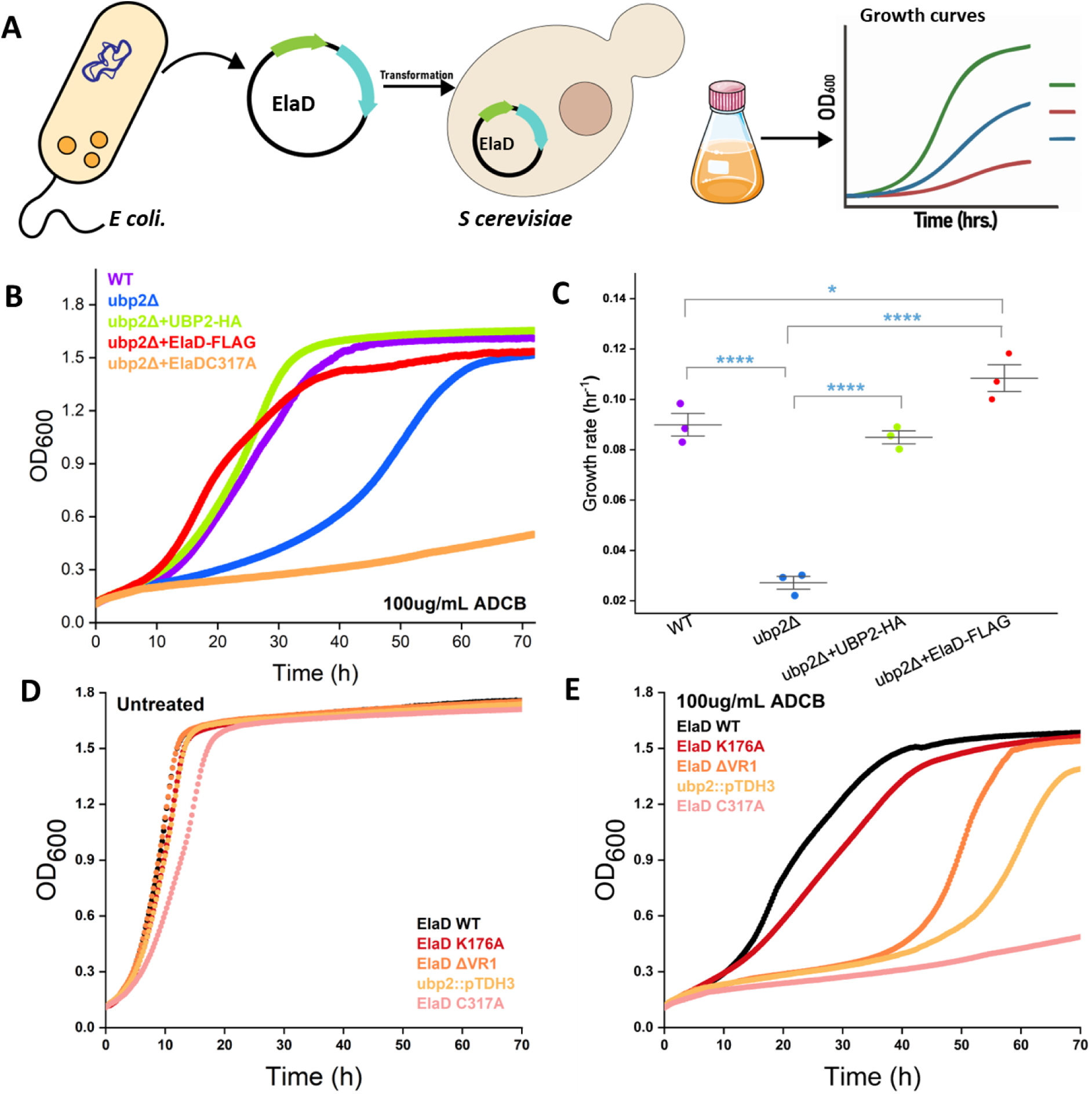
ElaD prevents the accumulation of K63-linked polyubiquitin chains in yeast under oxidative stress. A, Schematic representation of bacterial gene complementation in yeast. The gene encoding ElaD was cloned into the yeast integrative vector p406TDH3 (Addgene) and transformed into the *ubp2Δ* background. Growth kinetics were recorded under stress conditions, including H_2_O_2_ and the proteotoxic compound azetidine-2-carboxylic acid (ADCB), to evaluate phenotypic rescue. B, Growth curves for yeast strains with constitutive protein expression recorded on treatment with ADCB (100ug/mL). The *ubp2Δ* SUB280 strain complemented with Ubp2 was used as a positive control. C, Growth rates were calculated from the exponential phase of growth curves shown in Fig. 5B. Points represent independent biological replicates (n = 3), and bars indicate mean ± SD. Statistical significance was assessed using one-way ANOVA followed by Tukey’s multiple comparison test. *p < 0.05; ****p < 0.0001. D, Untreated yeast strains expressing different ElaD mutants show comparable growth in selective media. E, Growth curves for yeast strains treated with ADCB (100 µg/mL) show differential growth responses among ElaD mutants.

L-azetidine-2-carboxylic acid (ADCB) is a proline analogue known to induce widespread protein misfolding and disrupt cellular proteostasis in yeast [75,76]. Previous studies have shown that *ubp2Δ* yeast strains are hypersensitive to ADCB toxicity relative to wild-type cells, a phenotype linked to impaired regulation of K63-linked polyubiquitination and consequent defects in proteostasis [77,78]. The constitutive expression of Ubp2 is sufficient to rescue the phenotype in *ubp2Δ* yeast. We hypothesized ElaD could also rescue the yeast phenotype similar to Ubp2. Figure 5B shows a representative graph depicting constitutive expression of ElaD in the *ubp2Δ* yeast strain, which conferred a marked increase in resistance to the proteotoxic stressor. Growth assays demonstrated that ElaD-expressing *ubp2Δ* cells displayed significantly improved proliferation under ADCB treatment. To assess whether ElaD expression rescues the growth defect associated with loss of Ubp2, growth rates were compared across strains. A significant effect of genotype on growth rate was observed as u*bp2Δ* cells showed 4 times reduced growth relative to WT yeast. Expression of Ubp2 significantly restored growth compared with *ubp2Δ* and was indistinguishable from WT. Importantly, ElaD expression also robustly rescued the *ubp2Δ* growth defect, restoring growth rates close to WT levels, suggesting that ElaD can functionally compensate for loss of UBP2 activity in vivo (Figure 5C).

In contrast, yeast expressing the catalytically inactive ElaD mutant (ElaDC317A) exhibited pronounced growth defects compared to the *ubp2Δ* strain under identical conditions (Figure 5B). One possible explanation could be that the cellular resources are getting consumed in synthesizing this inactive protein, and the mutation leads to misfolding in the expressed protein. In support of the above explanation, we have observed that upon treatment with 0.6 mM H_2_O_2_, the *ubp2Δ ElaD* strain exhibited slower growth at later time points compared to the ubp2Δ strain (Figure S8), consistent with increased proteotoxic burden. Another possibility is increased sensitivity to ADCB-induced stress in the absence of functional ElaD and suggesting that the protective effect of ElaD against proteotoxic stress depends on its DUB activity. In this context, our findings suggest that ElaD can functionally compensate for the loss of Ubp2, likely by restoring K63-linked deubiquitination and thereby alleviating the accumulation of misfolded or stress-associated ubiquitinated proteins.

Having established this robust in vivo readout for ElaD activity, we leveraged the ADCB toxicity assay as a proof-of-concept to interrogate specific structural features required for K63-linked ubiquitin recognition in yeast. Based on the ElaD-Ub complex, we targeted two distinct structural features: the interfacial variable loop (ElaD ΔVR1) and a predicted electrostatic interaction (ElaD K176A). The variable region VR1 (aa 147-157) in ElaD also overlaps with the VR1 (aa 138-147) in SseL (Figure S9). Deletion of this unstructured loop in SseL abolished recognition of K63-linked diUb [52]. We engineered the mutant ElaD ΔVR1 strain to investigate the molecular basis of recognition of ubiquitin by ElaD. Another residue Lys176 in ElaD forms a H-bond contact with Glu51 in Ub (Figure 3C, top panel). As described previously, the DUB activity of wild-type ElaD rescues the *ubp2Δ* yeast phenotype from proteotoxic stress. Figure 5D shows similar growth in all strains under normal conditions. In contrast, when treated with 100ug/mL ADCB, the growth of cells expressing ElaDΔVR1 mutant was markedly compromised (Figure 5E). This suggests that the VR1 loop is critical for substrate recognition by ElaD. This impairment may reflect a requirement for this region in maintaining the proper fold of active conformation of ElaD or in mediating productive polyubiquitin engagement for the DUB activity. Conversely, the ElaD K176A mutant shows negligible effect on the phenotype. This result implies that other residues may be involved in the recognition of the substrate, or the substrate residues in fact play more important role for recognition.

### ElaD prevents hyperaccumulation of K63-linked polyubiquitin chains upon oxidative stress in yeast

To ascertain the functional role of ElaD in yeast, we investigated the accumulation of K63-linked polyubiquitination under oxidative stress in the ElaD C317A complemented strain. Under oxidative stress, increased K63-linked polyubiquitination of ribosomal proteins in yeast acts as a translation control mechanism that halts translation elongation [65]. As already mentioned, Ubp2, an endogenous DUB in yeast, is crucial to establish proteostasis for linked polyubiquitin from substrate proteins [63].

We hypothesize that ElaD should be able to prevent K63-linked polyubiquitin accumulation in yeast owing to its DUB activity and redox modulation. To evaluate this, a plasmid construct for constitutive expression using an auxotrophic selection was designed to express ElaD in *ubp2Δ* yeast. The accumulation of K63-linked polyubiquitin chains in response to oxidative stress was monitored in the subjecting yeast strains at different time points. Under oxidative stress conditions, yeast cells normally undergo a rapid and reversible translational arrest mediated by K63-linked polyubiquitination of ribosomes [20,21,61]. In oxidizing conditions, Ubp2 becomes inactive, resulting in the accumulation of K63-linked polyubiquitin chains and enforcement of translational arrest [79]. Consistent with previous reports, yeast cells lacking Ubp2 display elevated levels of polyubiquitination under oxidative stress [80]. We observed that the expression of ElaD in this background markedly reduced K63-linked polyubiquitination levels following H_2_O_2_ treatment for up to 2 h (Figure 6). Thus, ElaD expression alters the oxidative stress-induced ubiquitin signaling response in yeast. These results show that ElaD retains DUB activity under oxidative stress conditions and suppresses stress-induced K63-linked polyubiquitin accumulation in yeast.

**Figure 6.**
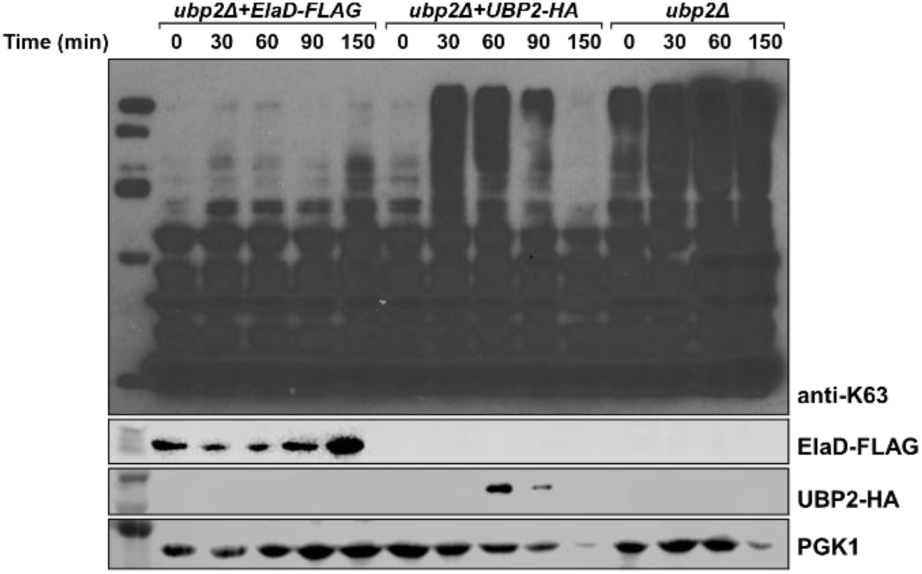
ElaD suppresses the accumulation of K63-linked polyubiquitin on exposure to oxidative stress. Immunoblot analysis of K63-linked polyubiquitin species from strains expressing ElaD-FLAG, Ubp2-HA, or no complementation. Cells were treated with 0.6 mM H_2_O_2_ for the indicated time points, and lysates were analyzed using anti-K63 ubiquitin antibody.

## DISCUSSION

This study identifies ElaD as a redox-regulated bacterial DUB that selectively recognizes ubiquitin and functionally rewires host ubiquitin-dependent stress responses. *In-vitro* biochemical assays demonstrate that ElaD activity is sensitive to the redox environment and undergoes reversible inhibition under oxidizing conditions, resembling regulatory mechanisms previously described for eukaryotic cysteine-based DUBs [26,32,81]. These findings extend the concept of redox control to CE-clan bacterial DUBs and suggest that bacterial effectors may exploit host oxidative conditions to fine-tune their enzymatic activity. In addition to redox-dependent regulation, our findings also provide insight into substrate recognition by ElaD. The results demonstrate that ElaD selectively recognizes Ub over Ubls, such as NEDD8, despite their structural similarity. Mutational analyses identified residues that contribute to modifier discrimination, and substitution of these residues altered substrate preference. These observations demonstrate that modifier selectivity in ElaD is governed by specific recognition determinants beyond the conserved catalytic core. Together, these findings support the broader view that substrate discrimination represents an important functional feature of bacterial CE-clan DUBs. Additionally, ElaD exhibits preferential activity toward K63-linked polyubiquitin chains, consistent with the linkage specificity reported for several bacterial DUBs involved in the modulation of host signaling pathways. We identified the structural features driving this discrimination and sought to clarify ElaD’s role within a physiological context. However, dissecting the specificity and physiological consequences of bacterial DUBs in mammalian systems remains challenging due to pathway redundancy and the complexity of innate immune networks [82–84]. By expressing ElaD in *S. cerevisiae* and monitoring changes in ubiquitin signaling under oxidative and proteotoxic stress conditions, we define how a pathogen-derived DUB interfaces with the physiologically relevant eukaryotic ubiquitin network. This approach provides mechanistic insights into how ElaD manipulates conserved stress signaling pathways.

*Saccharomyces cerevisiae* utilizes K63-linked polyubiquitination as a key regulatory response to oxidative stress, functioning as a non-proteasomal signal that enables adaptive translational control [80]. Endogenous DUB Ubp2 regulates this process through ROS-mediated inactivation, resulting in the accumulation of K63-linked polyubiquitin chains and enforcement of translational arrest, followed by subsequent recovery at later stages of stress adaptation. Interestingly, complementation of the ubp2Δ yeast strain with ElaD led to reduced accumulation of K63-linked polyubiquitin chains under oxidative stress conditions. The decrease in K63-linked polyubiquitination observed in ElaD-expressing yeast is consistent with perturbation of stress-associated translational regulation pathways previously linked to K63 ubiquitin signaling [85].ElaD-mediated removal of these ubiquitin chains may interfere with conserved adaptive stress responses associated with K63-linked ubiquitin signaling [32,66]. This effect might be due to the lack of regulatory pathways for ElaD, unlike eukaryotic DUBs that play a crucial role in the redox control of the translation pathway [63]. This interpretation is further supported by studies on other eukaryotic DUBs that show altered reactivity based on post-translational modifications [86,87], allosteric activation [88,89], and or interacting proteins [90]. Importantly, loss of K63-linked polyubiquitination does not substantially impair basal translation but instead disrupts stress-adaptive translational regulation, allowing continued elongation when translational arrest is required [20,65]. Consequently, proteostasis is compromised, and cellular sensitivity to oxidative stress is increased [75,76].

Functionally, ElaD suppresses ubiquitin-mediated proteostasis and stress signaling in yeast. We propose that ElaD-mediated suppression of K63-linked ubiquitination may alter adaptive stress-response pathways associated with translational regulation during oxidative stress. Additional support for this model comes from yeast growth assays under oxidative stress conditions, where ElaD expression resulted in reduced cell fitness. This growth defect may reflect altered regulation of oxidative stress adaptation pathways associated with K63-linked ubiquitin signaling. Although ElaD expression in yeast initially prevents stress-induced ubiquitin accumulation, the long-term consequence is compromised cellular adaptation, highlighting the regulatory nature of K63-linked ubiquitin signaling during oxidative stress. These findings suggest that bacterial DUBs may modulate host stress adaptation through premature removal of regulatory ubiquitin signals [91]. Together, these findings suggest that ElaD-mediated disruption of K63-linked ubiquitin signaling compromises stress adaptation and may represent a mechanism by which bacterial effectors interfere with host responses such as ROS generation during infection [37,92,93]. These results highlight how the bacterial DUB ElaD integrates redox resilience, substrate selectivity, and temporal control of ubiquitin signaling to modulate host pathways. In the context of infection, ElaD-mediated removal of stress-induced K63-linked ubiquitin chains might be important for delayed activation of host protective signaling pathways, thereby enabling pathogenic bacteria to transiently evade ubiquitin-dependent stress surveillance mechanisms. These principles may extend to other bacterial DUBs of the CE clan. More broadly, comparative studies of CE-clan DUBs will be important to determine whether redox sensitivity, selective substrate recognition, and targeting stress-adaptive ubiquitin signaling represent conserved strategies among bacterial effectors. Such analyses may reveal a shared mechanism by which CE-clan DUBs fine-tune host ubiquitin signaling in response to the dynamic redox environments encountered during infection.

## METHODS

### Cloning and protein expression

The plasmid encoding *elaD* gene was kindly provided by Dr. Jonathan Pruneda (Oregon Health & Science University, USA). To improve the solubility and recombinant expression of ElaD (elaD gene amplified by PCR from the BL21(DE3) *E. coli* strain), a plasmid construct with Silk-ElaD fusion sequence was generated. Full-length elaD gene was amplified by PCR for restriction site (*KpnI/XhoI*) insertion using primers:

Forward: 5’-GGAGGGTACCATGGTTACAGTTGTCAGCAATTATTGTCAATTATC-3’

Reverse: 5’-GGCACTCGAGTTAACTCACTCTTTTGCCGGATGC-3’

Existing pET26b+ vector in the laboratory, having the gene sequence for the mutant major ampullate spidroin 1 N-terminal domain (protein PDB ID: 6QJY), was digested with *KpnI/XhoI* for subsequent ligation with the elaD gene, resulting in Silk-ElaD chimera. Mutant ElaD constructs for yeast transformations were generated by site-directed mutagenesis following the manufacturer’s protocol using Q5 Hot Start High-Fidelity 2× Master Mix (New England Biolabs). To generate the di-ubiquitin substrate construct, a plasmid containing the coding sequences for both ubiquitin and NEDD8 was required. The ubiquitin plasmid was digested with *NheI* and *PstI* restriction enzymes to create compatible ends for NEDD8 insertion.

Plasmid containing the NEDD8 gene, pET28A/NEDD8, was obtained from Dr. Ranabir Das (NCBS, Bengaluru, India). NEDD8 was PCR-amplified using 2x PCR Master Mix (Thermo Scientific) and the following primers designed for *NheI/PstI* cloning:

Forward: 5′-GGTGCCGCGCGGCGCTAGCGGTGGCATGCTAATTAAAGTGAAG-3′

Reverse:5′-CGACGGAGCTCGAATTCCTGCAGGCCTCCTCCTCTCAGAGCCAACACC-3′

The amplified NEDD8 fragment was purified and ligated into the *NheI/PstI*-digested pET-Duet1/Ub plasmid commercially synthesized (Biotech Desk Pvt. Ltd.), using Anza™ Ligase Master Mix (Thermo Fisher Scientific). The resulting ligation mixture was transformed into *E. coli* DH5α cells, and positive clones were screened by digestion with *NheI* and *PstI*. Successful insertion of the NEDD8 gene was verified by agarose gel electrophoresis and confirmed by Sanger sequencing using the primers:

K48-Tail: 5′GCCGGTTAGCAGCTCGAGGACGGTAGAACGCTGT3′ and T7 terminator.

ΔElaD R2(DE3) cells were a kind gift from Prof. T. Hay, University of Dundee. Site-directed mutagenesis was performed using a ligase-independent method to create point mutations in the ubiquitin and NEDD8 sequences [94,95]. *E. coli* BL21-CodonPlus (DE3)-RIL cells were transformed with pOPINB/elaD, while E. coli BL21 and ΔElaD R2(DE3) cells were transformed with Silk-elaD and pETDuet1/Ub-NEDD8 plasmid constructs. A single transformed colony from each construct was used to inoculate 10 mL of LB medium and grown overnight at 37 °C with shaking at 200 rpm for 14–16 h. The overnight cultures were then used to inoculate 1 L of LB medium containing appropriate antibiotics, and the cells were grown at 37 °C until the optical density at 600 nm reached 1.0-1.2. This induction range was standardized for measurements on the Multiskan Go spectrophotometer (Thermo Fischer Scientific), as OD values may vary across instruments [96]. Protein expression was induced with 1 mM isopropyl-β-D-1-thiogalactopyranoside (IPTG), and cultures were incubated at 37°C for 4 h. Cells were harvested by centrifugation at 6000 rpm for 30-45 min at 4 °C, and the resulting pellets were stored at –80 °C until further use. For lysis, cell pellets were resuspended in lysis buffer (20 mM Tris-HCl, 200 mM NaCl, pH 7.4) supplemented with 1× protease inhibitor cocktail (Sigma-Aldrich) and disrupted by sonication on ice. The lysate was clarified by centrifugation at 17,000 rpm for 45 min at 4 °C, and the supernatant was subjected to Ni–NTA affinity chromatography.

The Ni–NTA resin was washed sequentially with lysis buffer and wash buffer containing 20 mM imidazole (in Tris-HCl, pH 7.7), followed by elution of the His_6_-tagged protein using 250 mM imidazole. The protein was further purified by size-exclusion chromatography using a Superdex 200 pg column (GE Healthcare) on a Bio-Rad Biologic Duo-Flow FPLC system. The purity and integrity of the eluted fractions were analyzed by SDS–PAGE, and the Ub-NEDD8 fusion protein was observed at the expected molecular weight (∼43.5 kDa) (Figure 4D).

### Yeast strains, plasmids, culture, and protein expression

All *Saccharomyces cerevisiae* strains and primers used in this study are described in Supplementary Table S5. Plasmids are listed in the supplementary table S6. pTDH3/ElaD was cloned using NEBuilder HiFi DNA Assembly Cloning Kit with restriction enzymes *XbaI* and *XhoI*. Yeast was transformed using the lithium acetate method [97]. Unless otherwise specified, yeast cells were cultivated in synthetic dextrose minimal medium (SD: 0.67% yeast nitrogen base, 2% dextrose, and required amino acids) at 30°C with shaking at 200 rpm. Cells were harvested at the exponential phase OD_600_ 0.3-0.5. Protein extraction and preparation for immunoblotting assays were performed as described previously [80].

### DUB activity assay

DUB activity was assessed using the fluorogenic substrates Ub–Rho (LifeSensors; excitation 485 nm, emission 535 nm). Reactions were performed in 50 mM Tris-HCl buffer (pH 7.5) containing 150 mM NaCl at 30 °C. Where indicated, samples were pre-incubated with peroxides or DTT for 5 minutes prior to initiating the reaction. For measurements of DUB activity in total cell extracts, cell lysis was performed in the absence of iodoacetamide (IAA) to preserve enzymatic activity.

### ESI-MS analysis

For electrospray ionization mass spectrometry (ESI-MS)-based analysis, the reaction mixtures were first buffer-exchanged using size exclusion chromatography on ENrich SEC 70 10 x 300 Column (Bio-Rad) into 20 mM ammonium acetate. Fractions corresponding to the protein peak were supplemented with 0.1% formic acid and analyzed by ESI-MS in positive ion mode.

### Immunoblotting

For confirmation of His-ElaD expression; immunoblots were performed using anti-His, anti-mouse IgG2a heavy chain (1:3000; cat. # ab97245). Accumulation of K63-linked ubiquitin chains under oxidative stress in yeast was assessed by immunoblotting with appropriate antibodies. Briefly, proteins extracted from cell lysate were quantified using the Bradford assay (Bio-Rad) before loading on the gel. Proteins were separated using 10% SDS-PAGE and transferred to a PVDF membrane for blotting. The antibodies used in this study were: anti-K63 ubiquitin (1:6,000, EMD Millipore, cat. # 05-1308, clone apu3), anti-ubiquitin (1:10,000; Cell Signaling Technology, cat. #3936S), anti-PGK1 (1:6,000; Invitrogen, cat. #22C5D8), anti-GAPDH (1:3,000, Abcam, cat. #ab9485), anti-HA (1:3,000, ThermoFisher, cat. #71-5500), anti-FLAG (1:3000, Sigma, cat. #F3165), anti-mouse IgG (1:3000, GeneTex, cat. # GTX221667-01) and anti-Rabbit IgG (1:4,000-10,000; Cytiva, cat. #NA934). Immunoblots were developed by chemiluminescence using the Amersham ECL Prime (Cytiva, cat. #RPN2232).

### Yeast growth curves

The sensitivity of yeast strains to the proteotoxic agent L-Azetidine-2-carboxylic acid (ADCB) was assessed. Strains were grown to exponential phase and then diluted to an OD600 of 0.1, after which growth was monitored over time. ubp2Δ SUB280 constitutively expressing ElaD also confers resistance to ADCB. Yeast strains expressing wild-type or mutant ElaD constructs were similarly monitored for growth at 30◦C and 200rpm upon treatment with ADCB. The yeast strain was complemented with different mutants of ElaD, and phenotype analysis was performed. Growth rates were calculated from the exponential phase of growth curves (11h-17h) by linear regression of ln(OD₆₀₀) versus time. Statistical comparisons between strains were performed using one-way analysis of variance (ANOVA) followed by Tukey’s multiple comparison test. Statistical significance was defined as p < 0.05. All analyses were performed using OriginPro (OriginLab).

### Complex prediction using ColabFold

The ElaD–Ubiquitin complex was predicted using ColabFold (AlphaFold2.ipynb; AlphaFold2-multimer v3 model parameters). The amino acid sequences of E. coli ElaD (UniProt: Q8XCY9) and ubiquitin (PDB: 1UBQ) were retrieved from the UniProt database. No signal peptides or non-native tags were included. ElaD and ubiquitin sequences were provided as separate chains using the separator “:” to enforce multimer prediction. Multiple sequence alignments (MSAs) were automatically generated by ColabFold using MMseqs2 against the UniRef and environmental sequence databases. Default model settings were used except that four independent seeds were generated to probe conformational variance in the predicted complex. Five unrelaxed models were generated per seed, and ranking was based on the internal AlphaFold pTM-weighted ranking score. Predicted confidence metrics were extracted from ranking files generated by AlphaFold. These included pLDDT, inter-chain predicted TM-score (ipTM), global pTM, and the predicted aligned error (PAE) matrix. Amber-relaxed models generated by ColabFold were used only for visualization and interaction mapping.

Structural alignment and RMSD analysis: For conformational consistency analysis, the top unrelaxed model from each seed was aligned to the seed-1 model in ChimeraX (version X.X). Root-mean-square deviation (RMSD) was computed using MatchMaker with pruning enabled. Two RMSDs were reported: pruned-atom RMSD (0.4-0.6 Å) reflecting the core interface after removal of outlier atoms, all-atom RMSD (1.4–2.8 Å) for the entire aligned region. The pruned value reflects interface similarity, whereas the full alignment captures global conformational differences.

Interface and SASA analysis: Interface residues were identified in ChimeraX using the “contacts” tool with a 5 Å heavy-atom distance cutoff. Solvent-accessible surface area (SASA) was computed using the measure “sasa” command with the default probe radius of 1.4 Å. Buried surface area (ΔSASA) was calculated as:

ΔSASA = SASA(A) + SASA(B) – SASA (A:B)
where A = ElaD, B = ubiquitin, and A:B = the complex. The resulting buried area (1609 Å²) was used as a measure of interface size. Hydrogen bonds and salt bridges were identified using ChimeraX H-bonds with default geometric criteria.

External validation tools: Binding affinity estimates were calculated using PRODIGY contacts [98–100]. Because PRODIGY overestimates affinity in predicted models, ΔG values were used qualitatively rather than quantitatively. The DockQ/pDockQ values were not used because AlphaFold2-multimer v3 already provides ipTM, which correlates strongly with the correct inter-chain orientation. All structural visualizations were performed in ChimeraX (UCSF). All computations were performed on Google Colab GPU instances running ColabFold. or ubiquitin over other ubiquitin-like proteins. To define residues critical for the ElaD-ubiquitin interaction, an ElaD–ubiquitin complex was modeled using AlphaFold3. Interface analysis revealed hydrogen-bonding interactions between the R72 and E51 ubiquitin residues and the proximal charged residues in ElaD. To experimentally validate these findings, site-directed mutagenesis was performed to generate diUb mutants: Ub R72A/ NEDD8 A72R and Ub E51N/ NEDD8 E51N.

### Supporting Information

Supplementary Table S2b displays all the contacts between ElaD and Ub. List of complemented yeast strains and primers used in the study is provided in Supplementary Table 5. List of protein sequences is provided in Supplementary Information.

## Supporting information

https://docs.google.com/document/d/1d2frDa6zgh0J4ceyEfl1-JEEsdTDp0ez/edit?usp=sharing&ouid=117501239392937517454&rtpof=true&sd=true

## Acknowledgements

S.R.K.A. acknowledges the financial support of the Department of Atomic Energy (DAE), India, under project no. 12-R&DTFR-5.10-0100. L.G. acknowledges the Fulbright-Nehru Doctoral Research Fellowship (Award No. 3045/FNDR/2024–2025) from the United States-India Educational Foundation (USIEF). L.G and S.R.K.A. acknowledge the support and resources provided by Duke University during the fellowship period. We thank the Silva Lab members at Duke University and SRKA lab members at TIFR, Mumbai, for valuable feedback and consistent support during this study.

## Author Contributions

L.G., S.R.K.A. and G.M.S conceived and designed the study. S.R.K.A. and G.M.S. supervised the work and provided laboratory resources. G.M.S. and S.R.K.A. contributed to experimental design and provided conceptual guidance. L.G. generated resources, performed experiments, and analyzed the data. A.S. helped in the design of the fusion protein construct and performed enzymatic assays. G.C.B. helped in designing the plasmids and yeast transformations. S.R.K.A., and L.G. wrote the manuscript. All authors contributed to its final form.

