## Supplementary material for "Redox-modulated bacterial deubiquitinase ElaD: Target recognition and suppression of K63-linked polyubiquitin accumulation in yeast": https://docs.google.com/document/d/1d2frDa6zgh0J4ceyEfl1-JEEsdTDp0ez/edit?usp=sharing&ouid=117501239392937517454&rtpof=true&sd=true

### Supplemental Figures

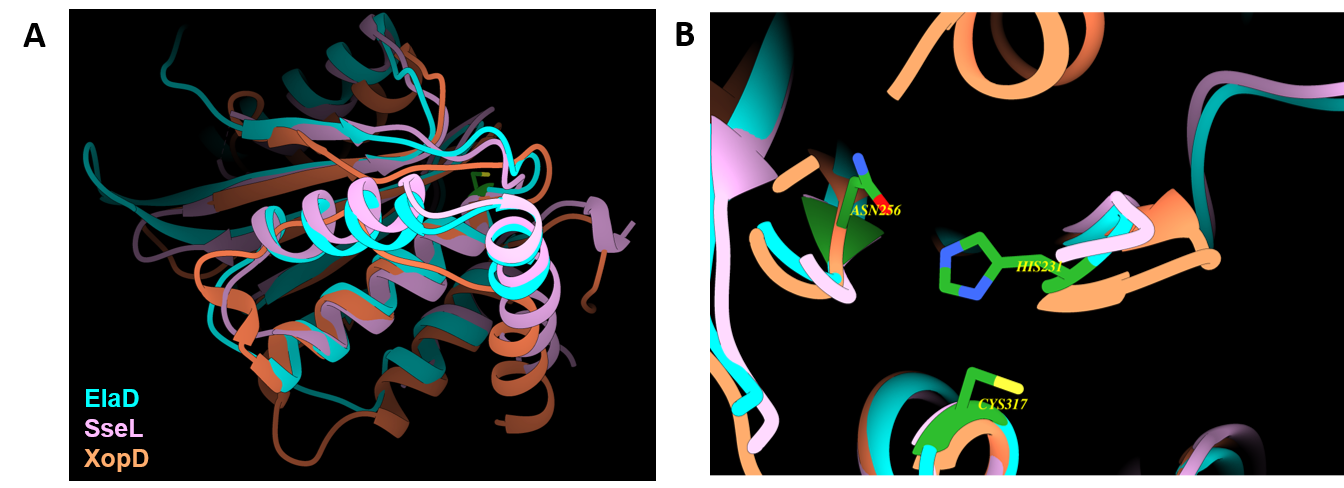

**Figure S1. Linkage specificity of bacterial cysteine proteases. Molecular graphics and structural analyses were performed with UCSF ChimeraX.**

### **A**, AlphaFold-predicted structure of ElaD, colored according to per-residue confidence scores (pLDDT). **B**, Structural overlay of the catalytic domains of ElaD and other indicated deubiquitinases (DUBs), shown in cartoon representation. Structural alignment was performed using UCSF Chimera. The catalytic triad of Cys317, His231 and Asn256 also overlaps with conserved residues in homologs.

**Figure S2. Confidence metrics for the predicted ElaD–ubiquitin complex.**
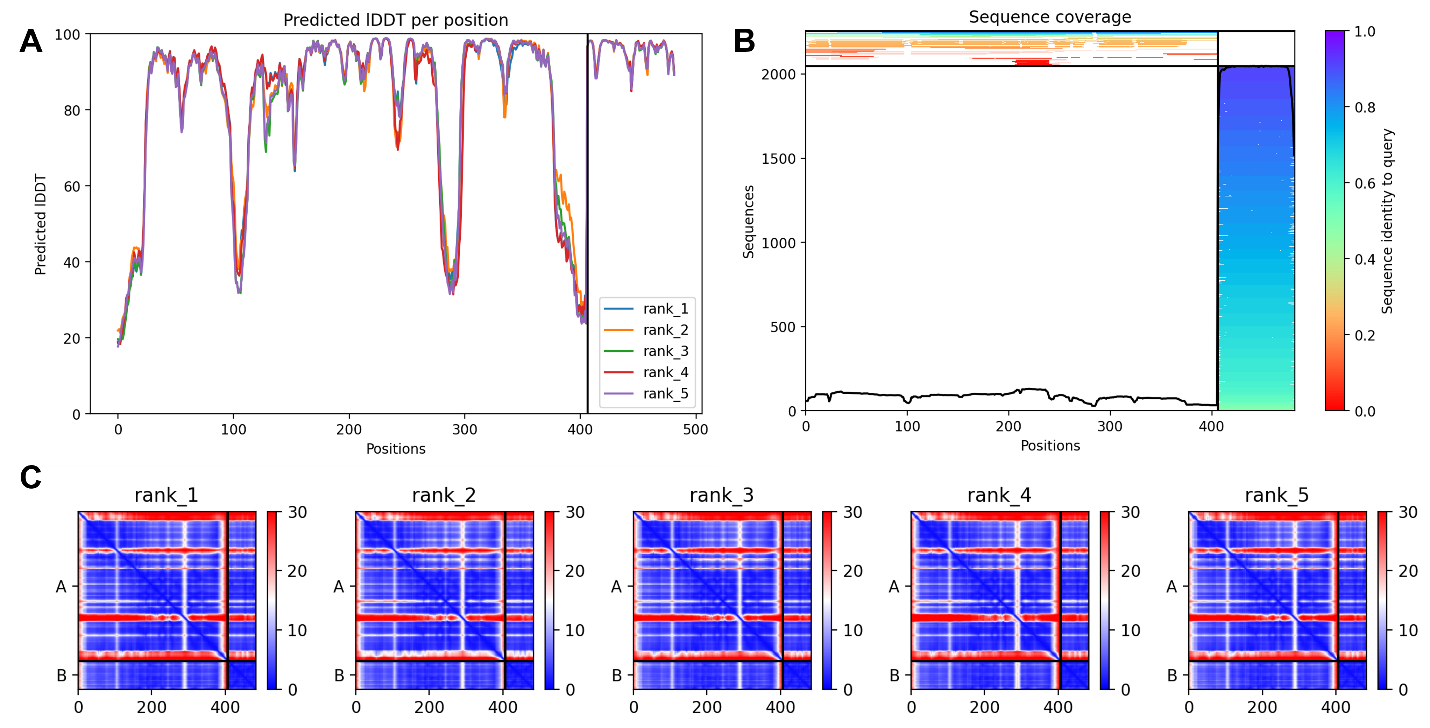

A, Predicted local distance difference test (pLDDT) scores for the highest-ranked model.

B, Sequence coverage of the ElaD and ubiquitin inputs used for structure prediction.

C, Predicted aligned error (PAE) plots for the five highest-ranked models, with emphasis on the protein–protein interaction interface.

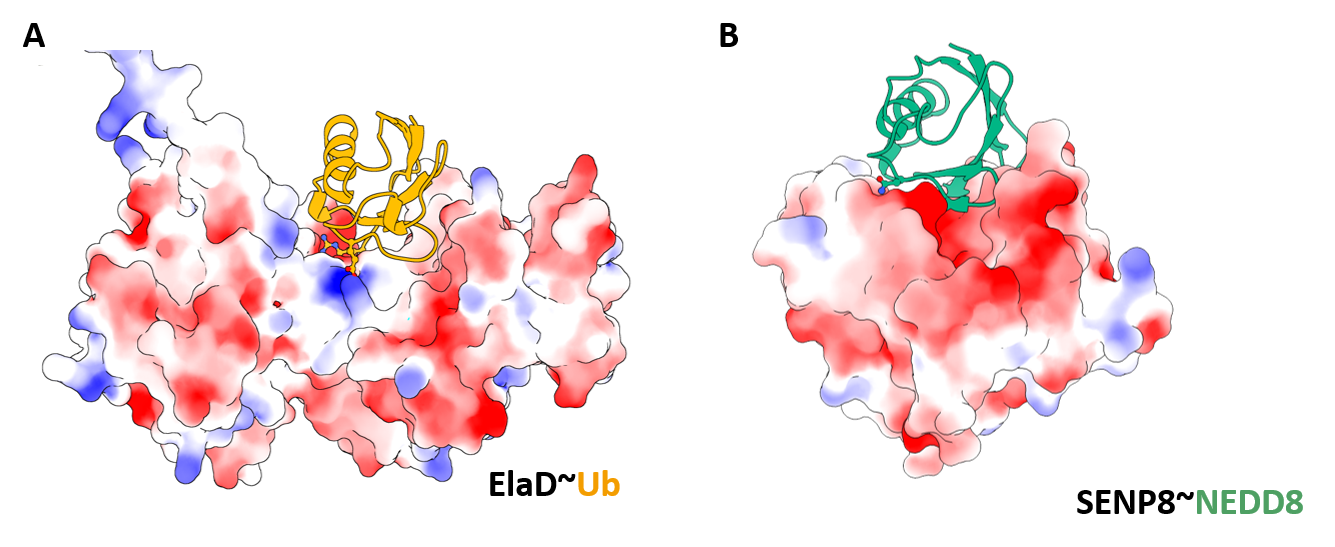

**Figure S3. Surface interactions drive substrate recognition**.

A, Cartoon representation of ElaD in complex with Ub highlighting complementary charged residues at the interaction interface.

B, Cartoon representation of SENP8 in complex with NEDD8 illustrating that interactions are electrostatically driven.

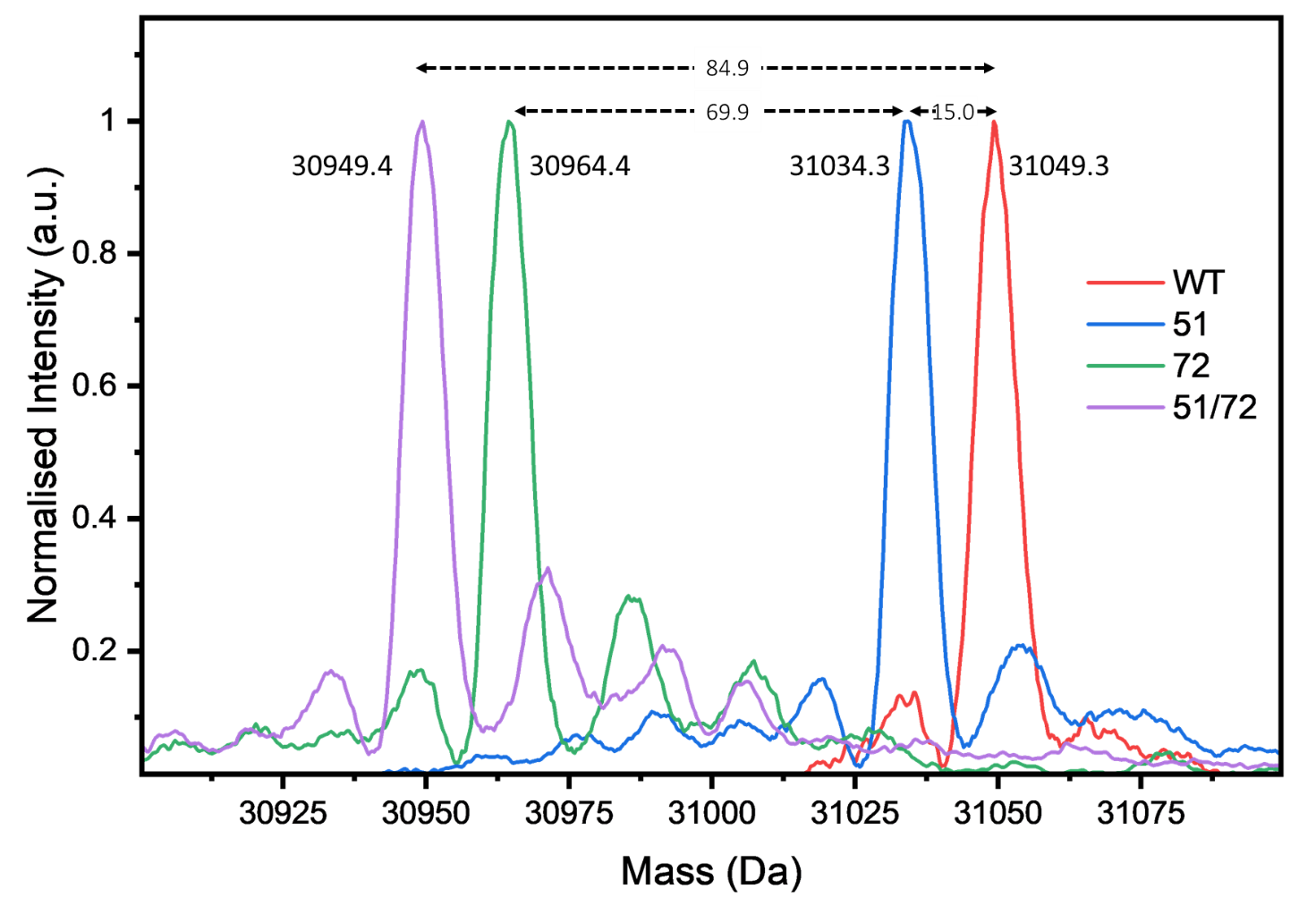

**Figure S4. ESI-MS analysis of Ub-NEDD8 substrate protein variants.**

Deconvoluted ESI-MS spectra of the purified (I27)_2_-Ub-containing fragment from the Ub-NEDD8 WT, 51, 72, and 51/72 substrate proteins. The observed masses match the expected changes introduced by the reciprocal substitutions: the 51 mutant showed a mass decrease of 15.0 Da, the 72 mutant showed a mass decrease of 69.9 Da, further from the 51 mutant, and the 51/72 double mutant showed the combined mass decrease of approximately 84.9 Da. These data confirm the expected molecular identities of the substrate variants used in this study.

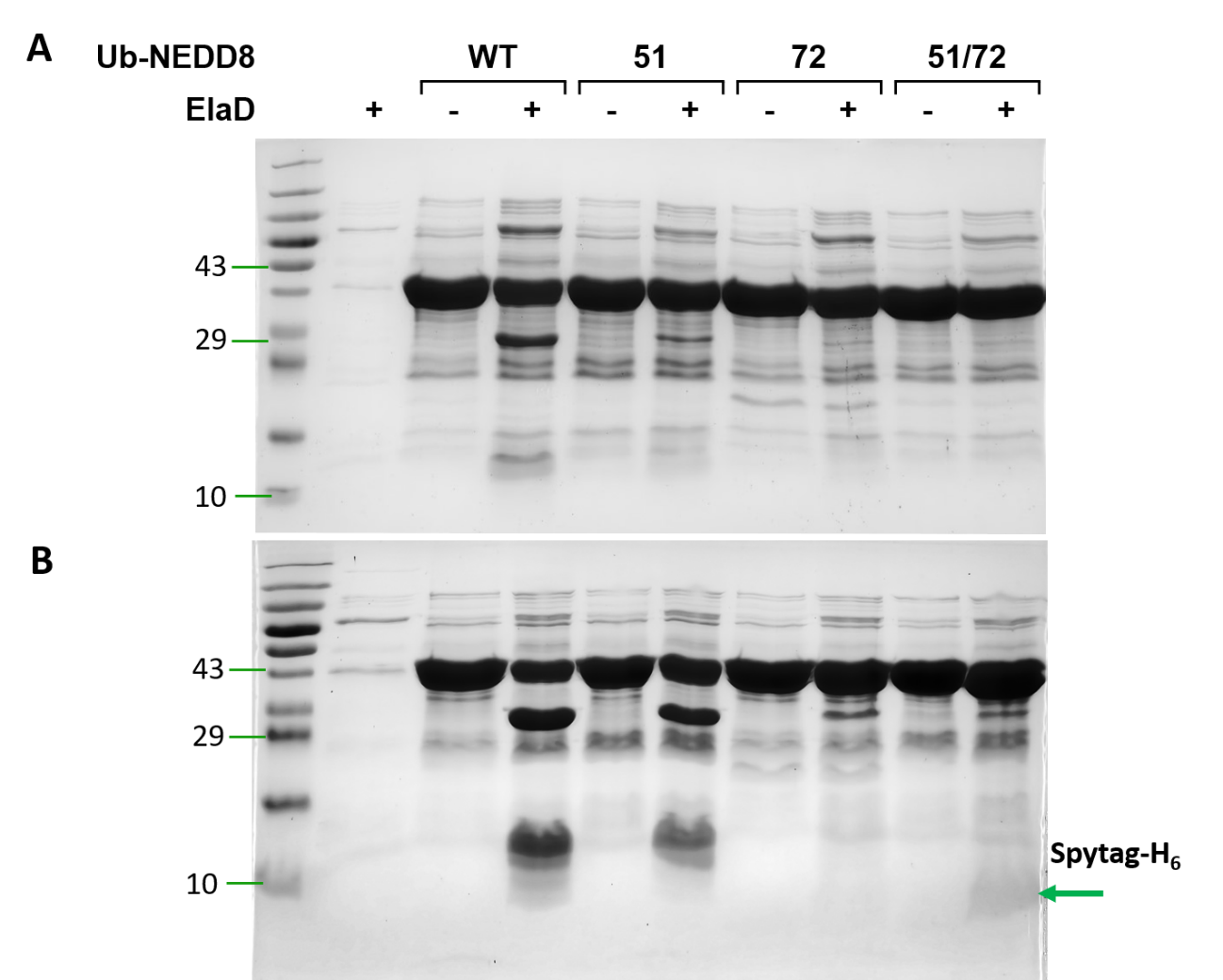

**Figure S5. ElaD shows similar cleavage patterns as Silk-ElaD, with slower endpoint cleavage.**
A, SDS-PAGE showing purified substrates cleavage after 30 min incubation with ElaD.
B, Same assay as A; reaction stopped after 4 h incubation.

The ElaD lane contains only the enzyme. WT denotes the unmodified Ub-NEDD8 substrate; 51 and 72 denote reciprocal substitutions at position 51 (UbE51N/N8N51E) and position 72 (UbR72A/N8A72R), respectively; and 51/72 denotes the combined double mutant. Paired lanes show substrates incubated in the absence (−) or presence (+) of ElaD. ElaD reproduced the same cleavage pattern observed with Silk-ElaD, although cleavage was weak at 30 min and became more apparent after 4 h. The red arrow shows the SpyTag-His_6_ fragment following cleavage at the NEDD8 (observed for the 51/72 mutant).

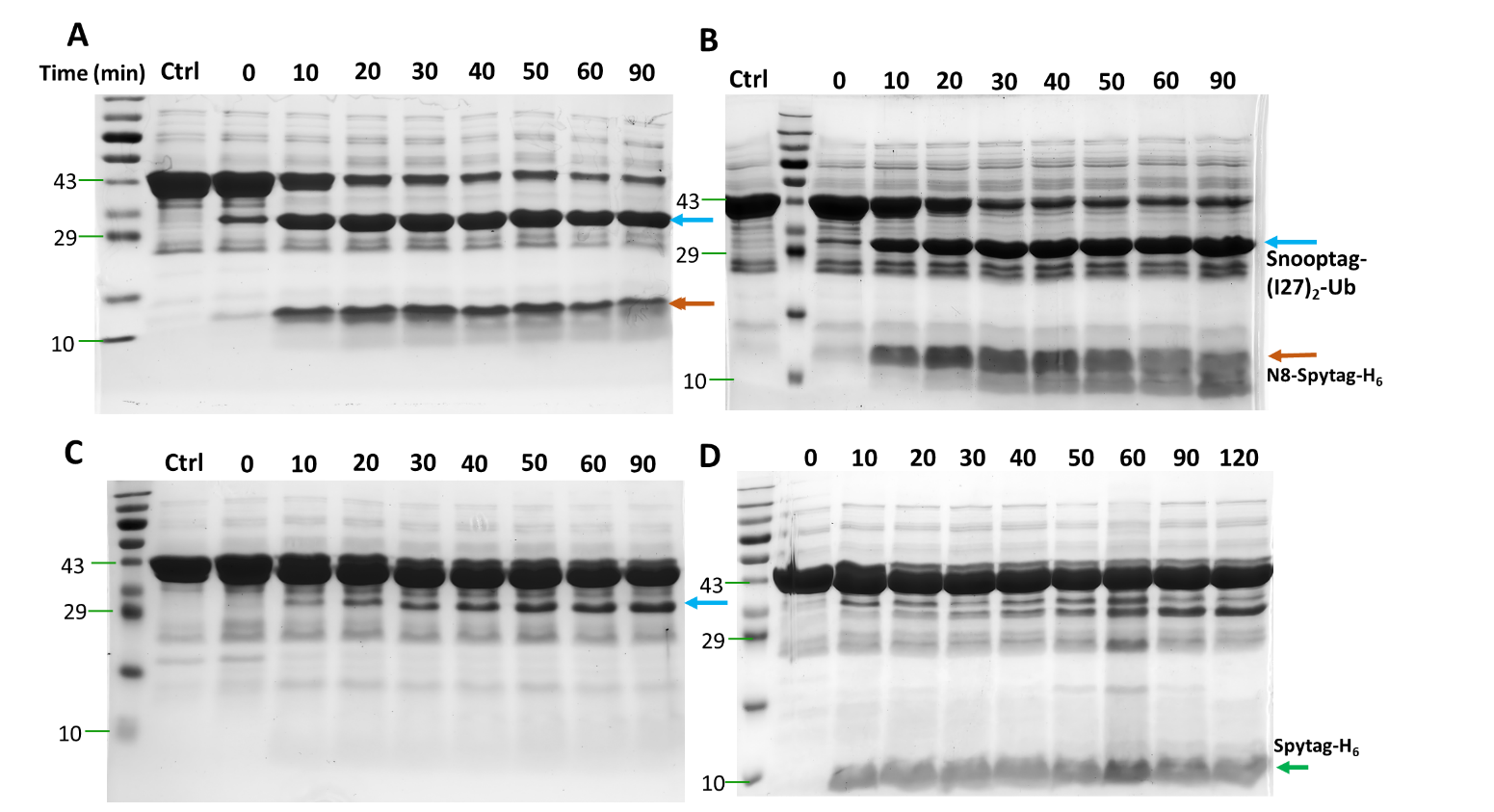

**Figure S6. Uncropped time-course SDS–PAGE used for analysis shown in Figure 4A.**

A, Ub-NEDD8 WT substrate treated with ElaD for indicated time points.

B, Ub-NEDD8 51 treated with ElaD for indicated time points.

C, Ub-NEDD8 72 substrate treated with ElaD for indicated time points.

D, Ub-NEDD8 51/72 double mutant substrate treated with ElaD for indicated time points.

Control denotes just the substrate. Reaction time points are indicated above the lanes.

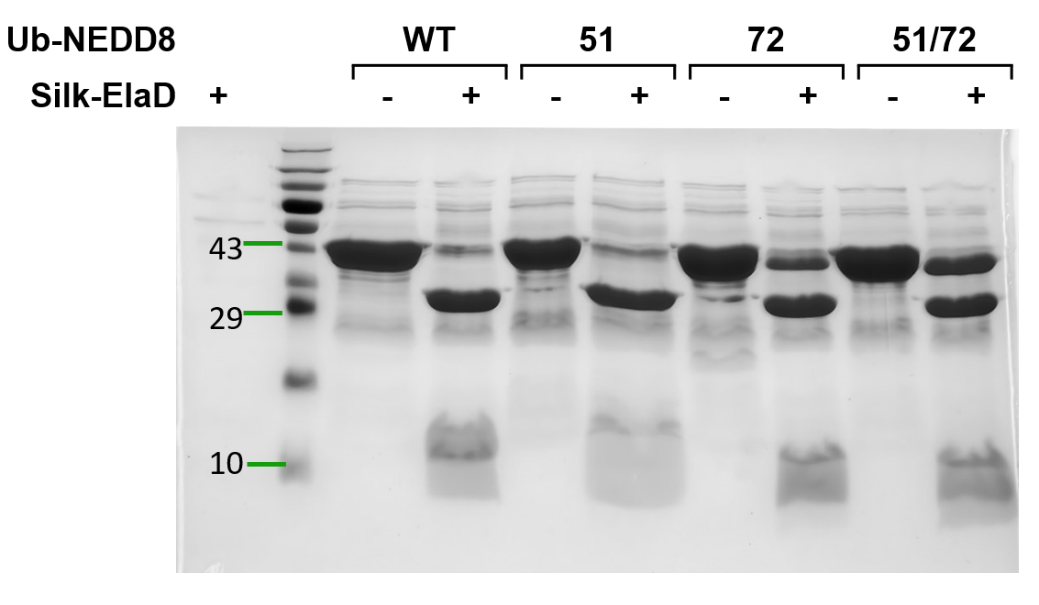

**Figure S7. 4 h endpoint assay reveals residual cleavage of the 72 and 51/72 substrates.**

Endpoint SDS-PAGE cleavage assay using Silk-ElaD after 4 h incubation.

Paired lanes show substrates incubated in the absence (−) or presence (+) of enzyme. WT denotes the unmodified Ub-NEDD8 substrate; 51 and 72 denote reciprocal substitutions at position 51 (UbE51N/N8N51E) and position 72 (UbR72A/N8A72R), respectively; and 51/72 denotes the combined double mutant. After 4 h incubation, both Ub-NEDD8 72 and Ub-NEDD8 51/72 mutant remained susceptible to cleavage, with detectable cleavage at the ubiquitin C-terminus. These data indicate that the mutations reduce and redirect cleavage rather than rendering the substrates completely resistant, consistent with altered substrate selectivity rather than absolute specificity.

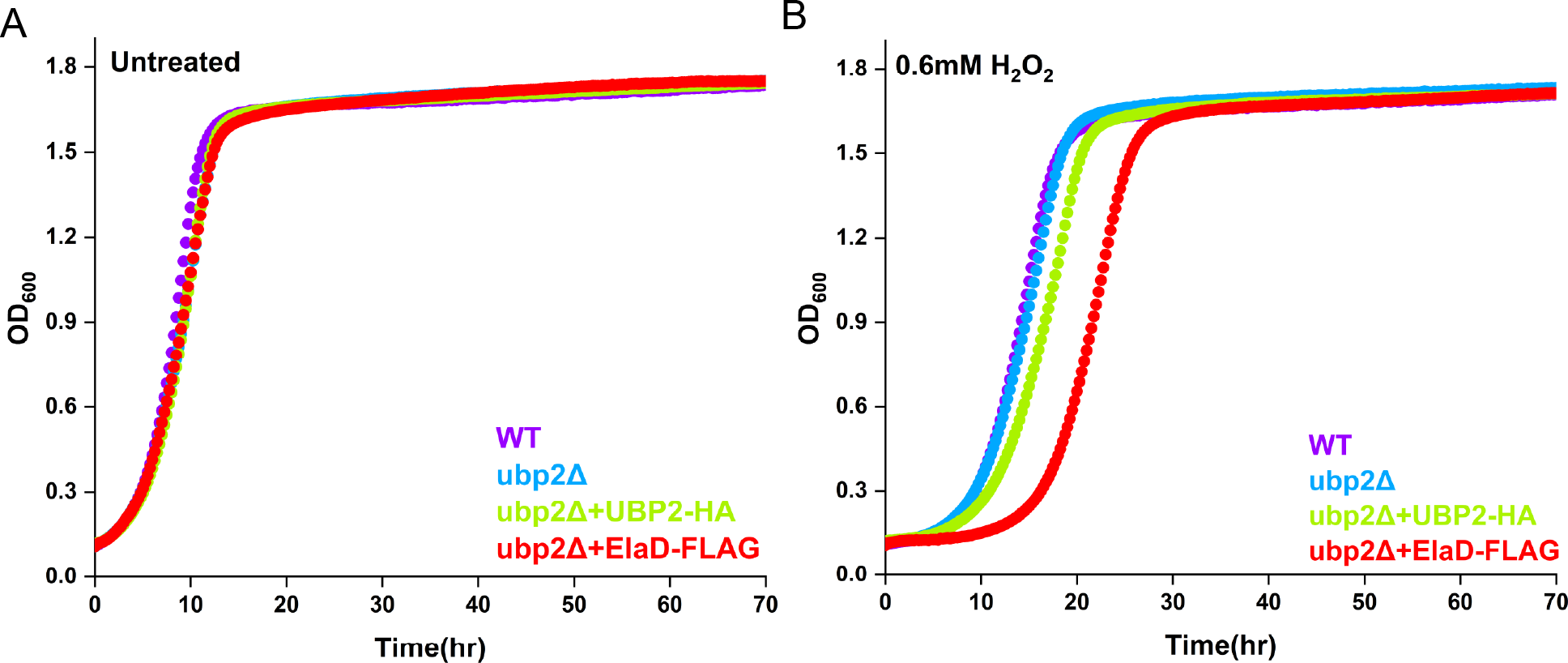

**Figure S8. Growth of yeast strains complemented with the indicated plasmids.**

A, The *Saccharomyces cerevisiae* SUB280 strain was used as the parental strain. The *ubp2Δ* mutant was used as available (1). For complementation, plasmids encoding either Ubp2 or the bacterial DUB ElaD under a constitutive promoter were introduced by transformation. Cultures were grown to mid-logarithmic phase and subsequently diluted to an OD_600_ of 0.1. Growth was monitored in appropriate selective media.

B, Cellular growth under oxidative stress was assessed by treatment with 0.6 mM H_2_O_2_. All strains were cultured in suitable media supplemented with H_2_O_2_.

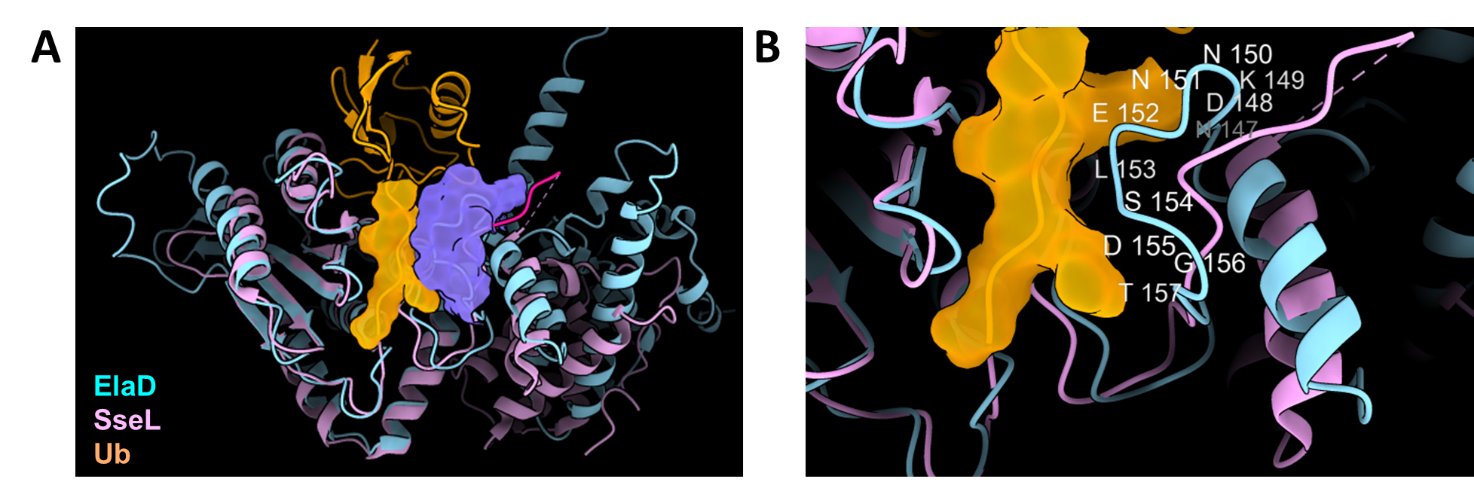

**Figure S9. Structural overlap of ElaD-Ub complex with SseL.**
A, The C-terminal region of ubiquitin forms contacts with the VR1 loop of ElaD, corresponding to an unstructured loop region in SseL.
B, VR1 loop is highlighted in the ElaD structure.

**Supplemental Tables**

**Table S1:** Pairwise structural deviations between the top-ranked model (#1) and selected alternative models. For each model, the overall rank and seed are indicated, along with pruned RMSD and full RMSD values for chain A and RMSD values for chain B. These measurements provide a quantitative assessment of structural similarity across independent docking solutions.

| Overall rank | Seed | Models | Pruned RMSD (chain A) | RMSD (Å) chain A | RMSD (Å) chain B |
| --- | --- | --- | --- | --- | --- |
| 1 | 2 |  |  |  |  |
| 3 | 3 | #2 to #1 | 0.568 | 1.378 | 0.192 |
| 4 | 1 | #3 to #1 | 0.405 | 2.775 | 0.042 |
| 8 | 0 | #4 to #1 | 0.362 | 2.15 | 0.037 |

**Table S2a**: List of intermolecular hydrogen bonds at the ElaD–ubiquitin interaction interface derived from structural modeling.

| Donor | Acceptor | Hydrogen | D..A Dist (Å) | D-H..A Dist (Å) |
| --- | --- | --- | --- | --- |
| /A ARG 388 NH1 | /B GLU 24 OE1 | /A ARG 388 HH12 | 2.594 | 1.793 |
| /A LYS 176 NZ | /B GLU 51 OE1 | /A LYS 176 HZ2 | 2.665 | 1.691 |
| /A GLN 310 NE2 | /B GLY 76 OXT | /A GLN 310 HE22 | 2.673 | 1.769 |
| /B ARG 42 NH2 | /A ASP 169 OD2 | /B ARG 42 HH21 | 2.724 | 1.748 |
| /B ARG 72 NE | /A GLU 152 OE2 | /B ARG 72 HE | 2.781 | 1.772 |
| /A TYR 189 N | /B GLY 47 O | /A TYR 189 H | 2.782 | 1.774 |
| /B ASN 60 ND2 | /A SER 213 OG | /B ASN 60 HD21 | 2.798 | 1.888 |
| /B LEU 73 N | /A THR 168 OG1 | /B LEU 73 H | 2.828 | 1.838 |
| /A THR 168 N | /B LEU 73 O | /A THR 168 H | 2.835 | 1.896 |
| /A ASN 210 ND2 | /B ALA 46 O | /A ASN 210 HD22 | 2.967 | 2.027 |
| /A ASN 227 ND2 | /B ARG 74 O | /A ASN 227 HD22 | 2.969 | 2.091 |
| /B GLY 76 N | /A GLY 230 O | /B GLY 76 H | 3 | 2.038 |
| /B ARG 72 NH1 | /A ASN 147 O | /B ARG 72 HH11 | 3.005 | 2.175 |
| /B GLN 40 NE2 | /A ASN 147 OD1 | /B GLN 40 HE21 | 3.023 | 2.445 |
| /A LYS 377 NZ | /B GLU 51 OE1 | /A LYS 377 HZ1 | 3.053 | 2.212 |
| /A CYS 317 N | /B GLY 76 OXT | /A CYS 317 H | 3.06 | 2.057 |
| /B ARG 72 N | /A GLU 152 OE2 | /B ARG 72 H | 3.062 | 2.112 |
| /A LYS 149 NZ | /B ASP 39 OD2 | /A LYS 149 HZ3 | 3.082 | 2.167 |
| /B GLN 40 NE2 | /A ASN 147 O | /B GLN 40 HE21 | 3.105 | 2.15 |
| /B ARG 72 NH2 | /A ASN 147 O | /B ARG 72 HH22 | 3.174 | 2.344 |
| /B ARG 42 NE | /A ASP 169 OD1 | /B ARG 42 HE | 3.283 | 2.28 |
| /B GLY 75 N | /A LEU 166 O | /B GLY 75 H | 3.401 | 2.549 |
| /A ASN 147 ND2 | /B ASP 39 OD2 | /A ASN 147 HD22 | 3.453 | 2.506 |
| /B ARG 72 NH2 | /A GLU 152 OE2 | /B ARG 72 HH21 | 3.484 | 2.713 |

**Table S3**: List of intermolecular hydrogen bonds at the USP2–ubiquitin interface.

| Donor | Acceptor | Hydrogen | D..A Dist (A˚) | D−H..A Dist (A˚) |
| --- | --- | --- | --- | --- |
| /B GLN 2 NE2 | /A THR 380 O | /B GLN 2 HE22 | 2.884 | 2.023 |
| /B LYS 6 NZ | /A PHE 367 O | /B LYS 6 HZ2 | 3.402 | 2.649 |
| /B LYS 6 NZ | /A ASP 371 OD2 | /B LYS 6 HZ3 | 2.843 | 1.889 |
| /B THR 14 N | /A GLU 377 OE1 | /B THR 14 H | 2.873 | 1.881 |
| /B ARG 42 NH1 | /A GLN 297 OE1 | /B ARG 42 HH12 | 2.877 | 2.043 |
| /B ARG 42 NH2 | /A GLN 297 OE1 | /B ARG 42 HH22 | 2.746 | 1.854 |
| /B GLN 49 NE2 | /A GLU 298 OE1 | /B GLN 49 HE22 | 2.832 | 1.882 |
| /B THR 66 N | /A ASP 345 OD1 | /B THR 66 H | 3.109 | 2.132 |
| /B ARG 72 NE | /A GLU 298 OE2 | /B ARG 72 HE | 2.774 | 1.873 |
| /B ARG 72 NH2 | /A GLU 298 OE1 | /B ARG 72 HH21 | 3.145 | 2.186 |
| /B LEU 73 N | /A ASP 295 OD2 | /B LEU 73 H | 2.699 | 1.693 |
| /B ARG 74 N | /A THR 460 O | /B ARG 74 H | 2.925 | 1.929 |
| /B ARG 74 NH1 | /A THR 460 O | /B ARG 74 HH11 | 2.964 | 2.128 |
| /B GLY 75 N | /A GLN 294 O | /B GLY 75 H | 2.951 | 2.110 |
| /B GLY 76 N | /A GLY 463 O | /B GLY 76 H | 3.175 | 2.200 |
| /A ASN 221 ND2 | /B GLY 76 OXT | /A ASN 221 HD21 | 3.146 | 2.333 |
| /A CYS 223 N | /B GLY 76 O | /A CYS 223 H | 3.214 | 2.322 |
| /A GLN 294 N | /B GLY 75 O | /A GLN 294 H | 2.978 | 1.990 |
| /A ARG 301 NE | /B GLY 47 O | /A ARG 301 HE | 2.765 | 1.881 |
| /A LYS 314A NZ | /B GLN 62 OE1 | /A LYS 314A HZ2 | 3.348 | 2.681 |

**Table S4**: List of intermolecular hydrogen bonds at the SENP8-NEDD8 interface.

| Donor | Acceptor | Hydrogen | D..A Dist (A˚) | D−H..A Dist (A˚) |
| --- | --- | --- | --- | --- |
| /B ARG 42 NE | /A ASP 29 OD1 | /B ARG 42 HE | 2.805 | 1.969 |
| /B ARG 42 NE | /A ASP 29 OD2 | /B ARG 42 HE | 3.077 | 2.198 |
| /B ARG 42 NH2 | /A ASP 29 OD2 | /B ARG 42 HH21 | 2.933 | 2.018 |
| /B LYS 48 NZ | /A ASP 10 OD2 | /B LYS 48 HZ3 | 3.022 | 2.229 |
| /B LYS 48 NZ | /A GLU 37 OE1 | /B LYS 48 HZ1 | 2.696 | 1.823 |
| /B ASN 51 ND2 | /A SER 7 OG | /B ASN 51 HD22 | 3.072 | 2.124 |
| /B ASN 51 ND2 | /A MET 9 O | /B ASN 51 HD21 | 2.912 | 1.915 |
| /B LYS 54 NZ | /A MET 9 O | /B LYS 54 HZ3 | 3.148 | 2.280 |
| /B TYR 59 OH | /A ASP 10 OD1 | /B TYR 59 HH | 2.702 | 1.790 |
| /B LEU 73 N | /A ASP 29 OD1 | /B LEU 73 H | 2.988 | 1.979 |
| /B ARG 74 N | /A GLY 99 O | /B ARG 74 H | 2.739 | 1.836 |
| /B GLY 75 N | /A LEU 27 O | /B GLY 75 H | 3.312 | 2.420 |
| /B GLY 76 N | /A THR 101 O | /B GLY 76 H | 3.149 | 2.157 |
| /A ASP 29 N | /B LEU 73 O | /A ASP 29 H | 2.930 | 2.131 |
| /A ASN 91 ND2 | /B ARG 74 O | /A ASN 91 HD22 | 3.007 | 2.026 |
| /A GLY 99 N | /B ALA 72 O | /A GLY 99 H | 2.687 | 1.777 |
| /A GLN 157 NE2 | /B GLY 76 O | /A GLN 157 HE22 | 3.283 | 2.469 |
| /A CYS 163 N | /B GLY 76 O | /A CYS 163 H | 2.612 | 1.692 |
| /B ARG 42 NE | /A ASP 29 OD1 | /B ARG 42 HE | 2.805 | 1.969 |
| /B ARG 42 NE | /A ASP 29 OD2 | /B ARG 42 HE | 3.077 | 2.198 |
| /B ARG 42 NH2 | /A ASP 29 OD2 | /B ARG 42 HH21 | 2.933 | 2.018 |
| /B LYS 48 NZ | /A ASP 10 OD2 | /B LYS 48 HZ3 | 3.022 | 2.229 |
| /B LYS 48 NZ | /A GLU 37 OE1 | /B LYS 48 HZ1 | 2.696 | 1.823 |
| /B ASN 51 ND2 | /A SER 7 OG | /B ASN 51 HD22 | 3.072 | 2.124 |

**Table S6:** List of plasmids used in the study.

| **Vector** | **Gene** | **Mutation** | **Tag** | **Protein** | **Reference** |
| --- | --- | --- | --- | --- | --- |
| pYES 2.0 | UBP2 | - | HA (C-term) | Ubp2-HA | *Clara et al., 2024* |
| pTDH3 | elaD | - | FLAG(C-term) | ElaD-FLAG | *this study* |
| pOPINB | elaD | - | His_6_ (N-term) | His-ElaD | *Pruneda et al., 2016* |
| pOPINB | ElaD C317A | C317A | His_6_ (N-term) | ElaD C317A | *this study* |
| pET2b+ | Silk-elaD | - | His_6_ (N-term) | Silk-ElaD | *this study* |
| pETDuet1 | Snooptag-(I27)_2_-Ub-NEDD8-Spytag-His_6_ | - | His_6_ (C-term) | Ub-NEDD8 WT | *this study* |
| pETDuet1 | Snooptag-(I27)_2_-Ub-NEDD8-Spytag-His_6_ | Ub E51N/  N8 N51E | His_6_ (C-term) | Ub-NEDD8 51 | *this study* |
| pETDuet1 | Snooptag-(I27)_2_-Ub-NEDD8-Spytag-His_6_ | Ub R72A /  N8 A72R | His_6_ (C-term) | Ub-NEDD8 72 | *this study* |
| pETDuet1 | Snooptag-(I27)_2_-Ub-NEDD8-Spytag-His_6_ | Ub E51N, R72A / N8 N51E, A72R | His_6_ (C-term) | Ub-NEDD8 51/72 | *this study* |
| pTDH3 | ElaD WT | - | FLAG(C-term) | - | *this study* |
| pTDH3 | ElaD C317A | C317A | FLAG(C-term) | - | *this study* |
| pTDH3 | ElaD ΔVR1 | ΔVR1 | FLAG(C-term) | - | *this study* |
| pTDH3 | ElaD K176A | K176A | FLAG(C-term) | - | *this study* |

**Protein Sequences**

SnoopTag-(I27)₂-Ub-N8-SpyTag-His_6_

1. Ub-NEDD8 WT

MGKLGDIEFIKVNKGSGEGGGSLIEVEKPLYGVEVFVGETAHFEIELSEPDVHGQWKLKGQPLAASPDCEIIEDGKKHILILHNCQLGMTGEVSFQAANTKSAANLKVKELRSLGLIEVEKPLYGVEVFVGETAHFEIELSEPDVHGQWKLKGQPLAASPDCEIIEDGKKHILILHNCQLGMTGEVSFQAANTKSAANLKVKELGRGELMQIFVKTLTGKTITLEVEPSDTIENVKAKIQDKEGIPPDQQRLIFAGKQLEDGRTLSDYNIQKESTLHLVLRLRGGASGGMLIKVKTLTGKEIEIDIEPTDKVERIKERVEEKEGIPPQQQRLIYSGKQMNDEKTAADYKILGGSVLHLVLALRGGLQGGSGRGVPHIVMVDAYKRYKGSGHHHHHH

2. Ub-NEDD8 51 (Ub E51N/ N8 N51E)

MGKLGDIEFIKVNKGSGEGGGSLIEVEKPLYGVEVFVGETAHFEIELSEPDVHGQWKLKGQPLAASPDCEIIEDGKKHILILHNCQLGMTGEVSFQAANTKSAANLKVKELRSLGLIEVEKPLYGVEVFVGETAHFEIELSEPDVHGQWKLKGQPLAASPDCEIIEDGKKHILILHNCQLGMTGEVSFQAANTKSAANLKVKELGRGELMQIFVKTLTGKTITLEVEPSDTIENVKAKIQDKEGIPPDQQRLIFAGKQLNDGRTLSDYNIQKESTLHLVLRLRGGASGGMLIKVKTLTGKEIEIDIEPTDKVERIKERVEEKEGIPPQQQRLIYSGKQMEDEKTAADYKILGGSVLHLVLALRGGLQGGSGRGVPHIVMVDAYKRYKGSGHHHHHH

3. Ub-NEDD8 72 (Ub R72A / N8 A72R)

MGKLGDIEFIKVNKGSGEGGGSLIEVEKPLYGVEVFVGETAHFEIELSEPDVHGQWKLKGQPLAASPDCEIIEDGKKHILILHNCQLGMTGEVSFQAANTKSAANLKVKELRSLGLIEVEKPLYGVEVFVGETAHFEIELSEPDVHGQWKLKGQPLAASPDCEIIEDGKKHILILHNCQLGMTGEVSFQAANTKSAANLKVKELGRGELMQIFVKTLTGKTITLEVEPSDTIENVKAKIQDKEGIPPDQQRLIFAGKQLEDGRTLSDYNIQKESTLHLVLALRGGASGGMLIKVKTLTGKEIEIDIEPTDKVERIKERVEEKEGIPPQQQRLIYSGKQMNDEKTAADYKILGGSVLHLVLRLRGGLQGGSGRGVPHIVMVDAYKRYKGSGHHHHHH

4. Ub-NEDD8 51/72 (Ub E51N, R72A / N8 N51E, A72R)

MGKLGDIEFIKVNKGSGEGGGSLIEVEKPLYGVEVFVGETAHFEIELSEPDVHGQWKLKGQPLAASPDCEIIEDGKKHILILHNCQLGMTGEVSFQAANTKSAANLKVKELRSLGLIEVEKPLYGVEVFVGETAHFEIELSEPDVHGQWKLKGQPLAASPDCEIIEDGKKHILILHNCQLGMTGEVSFQAANTKSAANLKVKELGRGELMQIFVKTLTGKTITLEVEPSDTIENVKAKIQDKEGIPPDQQRLIFAGKQLNDGRTLSDYNIQKESTLHLVLALRGGASGGMLIKVKTLTGKEIEIDIEPTDKVERIKERVEEKEGIPPQQQRLIYSGKQMEDEKTAADYKILGGSVLHLVLRLRGGLQGGSGRGVPHIVMVDAYKRYKGSGHHHHHH

5. His_6-_ElaD

HHHHHHSSGLEVLFQGPMVTVVSNYCQLSQTQLSQTFAEKFTVTEELLQSLKKTALSGDEESIELLHNIALGYDEFGKKAEDILYHIVRNPTNDTLSIIKLIKNACLKLYNLAHTATKHPLKSHDSDNLLFKKLFSPSKLMAIIGEDIPLISEKQSLSKVLLNDKNNELSDGTNFWDKNRQLTTDEIACYLKKIAANAKNTQVNYPTDFYLPNSNSTYLEVALNDNIKSDPSWPKEVQLFPINTGGHWILVSLQKIVNEKNNTQQIKCIIFNSLRALGHEKENSLKRIINSFNSFNCDPTRETPNNKNITDHLTEPEIIFLHADLQQYLSQSCGAFVCMAAQEVIEQMESNSDSAPYTLLKNYADRFKKYSAEEQYEIDFQHRLENRNCYLDKYGDANINHYYRNLEIKNSHPKNRASSKRVS

6. His_6_-Silk-ElaD

HHHHHHSHTTPWTNPGLAENFMNSFMQGLSSMPGFTASQLDKMSTIAQSMVQSIQSLAAQGRTSPNDLQALNMAFASSMAEIAASEEGGGSLSTKTSSIASAMSNAFLQTTGVVNQPFINEITQLVSMFAQAGMNDVSAENLYFQGGTMVTVVSNYCQLSQTQLSQTFAEKFTVTEELLQSLKKTALSGDEESIELLHNIALGYDKFGKEAEDILYHIVRTPTNETLSIIRLIKNACLKLYNLAHIATNSPLKSHDSDDLLFKKLFSPSKLMTIIGDEIPLISEKQSLSKVLLNDENNELSDGTNFWDKNRQLTTDEIACYLQKIAANAKNTQVNYPTGLYVPYSTRTHLEDALNENIKSDPSWPNEVQLFPINTGGHWILVSLQKIVNKKNNKLQIKCVIFNSLRALGYDKENSLKRVINSFNSELMGEMSNNNIKVHLNEPEIIFLHADLQQYLSQSCGAFVCMAAQEVIEQRESNSDSAPYTLLKNYADRFKKYSAEEQYEIDFQHRLANRNCYLDKYGDANINHYYRNLEIKHSQPKNRASGKRVS
